# Sub-Nanomolar Detection and Discrimination of Microcystin Congeners Using Aerolysin Nanopores

**DOI:** 10.1101/2025.07.15.664886

**Authors:** Alissa Agerova, Juan Francisco Bada Juarez, Luciano A. Abriata, Maria J. Marcaida, Anna Carratalà, Elizabeth M. L. Janssen, Chan Cao, Tamar Kohn, Matteo Dal Peraro

**Affiliations:** Institute of Bioengineering, School of Life Science, École Polytechnique Fédérale de Lausanne (EPFL), 1015 Lausanne, Switzerland; Laboratory of Environmental Virology, School of Architecture, Civil and Environmental Engineering (ENAC), École Polytechnique Fédérale de Lausanne (EPFL), Switzerland; Department of Environmental Chemistry, Swiss Federal Institute of Aquatic Science and Technology (EAWAG), Dübendorf 8600, Switzerland; Department of Inorganic and Analytical Chemistry, School of Chemistry and Biochemistry, University of Geneva, Geneva, CH-1205

## Abstract

Climate-driven disruptions in aquatic ecosystems are amplifying cyanotoxin production, threatening drinking and recreational water safety. Monitoring of these toxins is challenged by requirements of the low µg/L detections limits and structural diversity. Here, we employ aerolysin nanopores to distinguish seven of the most prevalent microcystin congeners, both individually and in mixtures, at environmentally relevant concentrations. Importantly, we showed that aerolysin enables the detection of microcystins in different lake water samples at concentrations below the World Health Organization’s intervention thresholds, reaching picomolar sensitivity. Moreover, combining experiments and molecular dynamics simulations, we further investigated the microcystin sensing mechanism, suggesting that the ionic current blockage is primarily governed by K238 in aerolysin, while dwell time is regulated by R220 constriction site. Our results open the way to use the nanopore sensing technology for real-time monitoring of microcystins in drinking water sources and surface waters.

## Introduction

Aquatic ecosystems and water quality are increasingly threatened by both anthropogenic pollutants and natural toxins, particularly those produced by cyanobacteria^1^. Climate change, eutrophication, and rising CO₂ levels have led to more frequent cyanobacterial blooms^2,3,4^, which negatively impact biodiversity^2^ and pose significant risks to human health^5^. During these blooms, cyanobacteria release a range of potent cyanotoxins — including neurotoxins, cytotoxins, and hepatotoxins such as microcystins, nodularins, and cylindrospermopsin — that inhibit key metabolic enzymes^6–9^. Cyanotoxins, especially microcystins, can accumulate to micromolar concentrations in surface scums^10^ and nanomolar levels in the water column^11^, well above the 1 µg/L safety guideline for chronic exposure to drinking water set by the World Health Organization (WHO)^12^. Even at lower concentrations, these toxins may persist before and after blooms in meso-oligotrophic lakes like Lake Geneva (Switzerland)^13^, impacting aquatic life despite being below regulatory thresholds.

Microcystins (MCs)^14,15^ are cyclic peptides made of canonical and three non-canonical amino acids: D-erythro-β-methyl-isoaspartic acid (D-Masp), 3-amino-9-methoxy-2,6,8-trimethyl-10-phenyldeca-4,6-dienoic acid (Adda), and N-methyldehydroalanine (Mdha) for the congener MC-LR (**Fig. 1A**, located at positions 3, 5, 7, respectively). Variations of this general scaffold can produce more than 300 congeners identified to date^16,17^. MCs are well-documented public health hazards; for example, a 1996 cyanobacterial bloom in Brazil led to the deaths of over 50 dialysis patients due to acute liver failure^18,19^. This incident highlighted the need for effective monitoring of MC concentrations in water. Human exposure typically occurs via oral or respiratory routes^19^, with symptoms ranging from mild skin rashes and nausea to severe gastrointestinal, liver damage and respiratory distress. At the molecular level, MCs are actively taken up by hepatocytes and potently inhibit the protein phosphatases PP1 and PP2A, disrupting cellular signaling and cytoskeletal integrity^20,21^, which can lead to cell death and, in severe cases, fatal liver failure^22^. The use of contaminated water for irrigation also raises concerns for food safety^23^. In response to these risks, the WHO established three guideline values **(Table S1)** that trigger specific interventions^12^, with the guideline for drinking water set at 1 µg/L (equivalent to 1 nM) for the best-studied microcystin congener, MC-LR^23^.

**Figure 1.**
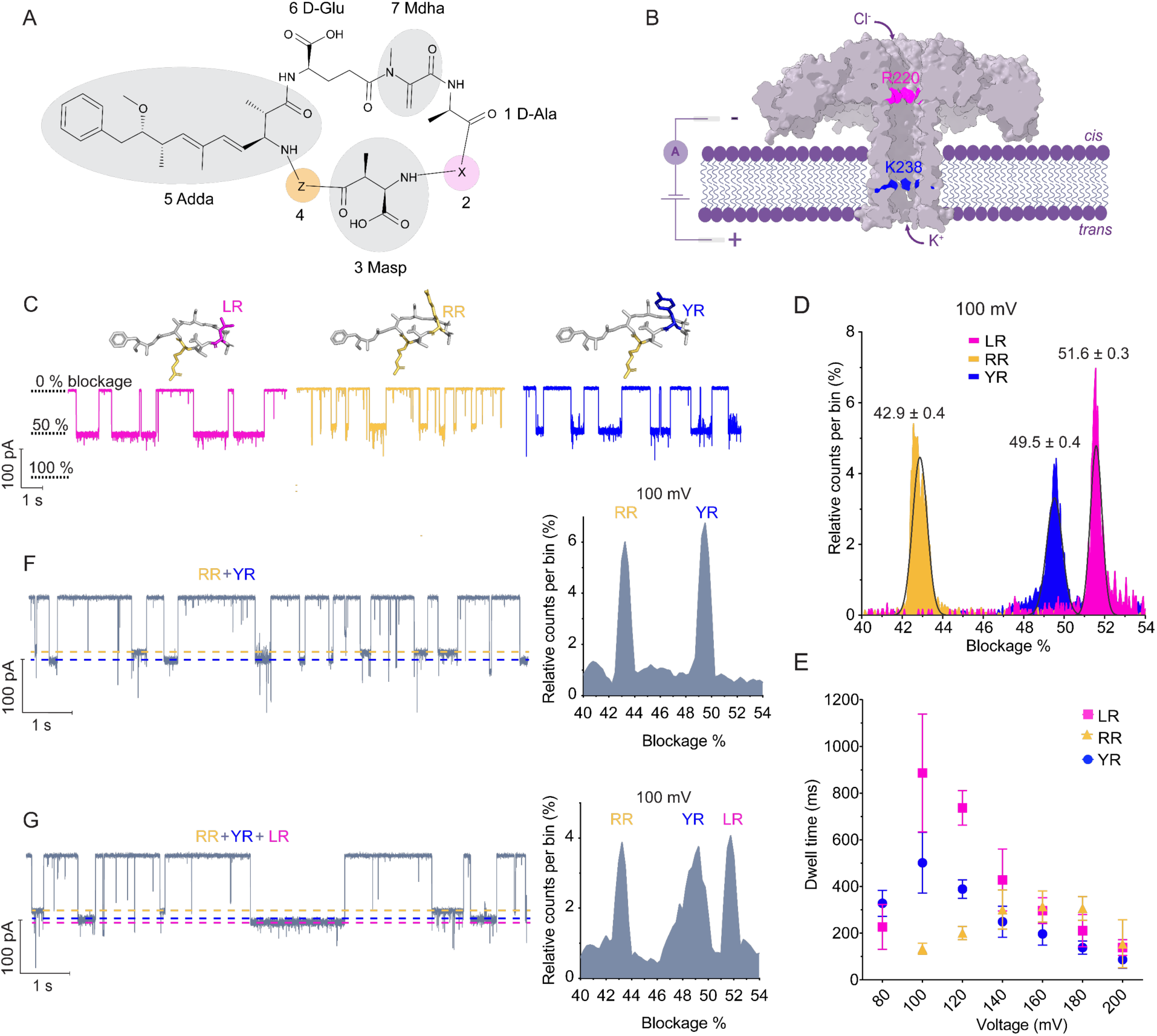
**Detection of MC-LR, MC-RR, MC-YR with WT AeL nanopore**. (**A)** Microcystin 2D structure with its non-canonical residues (gray), and 2 variable regions: X (pink), Z (yellow). **(B)** Cartoon of the WT AeL pore with two narrowest constrictions: R220 (magenta), K238 (blue). Membrane (purple), MC addition to the *cis* chamber (not to scale), passage of potassium and chloride ions are shown. **(C)** Typical traces recorded at 100 mV in 4 M KCl buffered with 10 mM Tris, 1.0 mM EDTA at pH 7.5. Variations in blockage signatures can be distinguished among the 3 congeners. Selected 10 s fragments of the current traces, recorded at a sampling rate of 50 kHz and filtered with a 500 Hz low-pass filter, are shown. **(D)** Histograms of current blockage produced by MCs at 100 mV in individual experiments, showing the current separation between the three different MCs. **(E)** Dwell time dependence on voltage. Each point is the result of the average of at least 3 pore replicates (**Figs. S1-S3, S5-S8**). **(F, G)** Raw current traces (recorded at 50 kHz, filtered to 500 Hz) and histograms (bin size = 0.25) showing blockage events of MCs and their distribution in a mixture of either two or three congeners.

To date, MCs are commonly monitored by enzyme-linked immunosorbent assay (ELISA, US EPA Method 546^24^), which is easy to operate and enables a rapid, picomolar-range^25^ detection. However, ELISA lacks specificity for individual MC congeners^26,27^ and may cross-react with similar toxins^28,29^, potentially causing false positives. In contrast, liquid chromatography coupled to mass spectrometry (LC-MS/MS, US EPA Method 544^30^) delivers the precise, sensitive and selective analysis of 11 MC congeners for which reference standards are available^30^, reaching sensitivities in the low nanomolar range (compatible with environmental samples^31^). However, the high cost of equipment, the need for qualified users, sample transportation and the long turnaround time until results are available make this method expensive and impractical for real-time analysis^30^. Currently, LC-MS/MS analysis of MCs takes from one day for quantitative results to several weeks for detailed suspect screening where no reference standards are available^32^. This delay limits timely interventions and increases the risk of missing sudden toxin spikes, as cyanotoxin levels can rapidly exceed safe limits due to blooms or cell lysis^33^. Real-time monitoring is therefore essential for promptly detecting toxin peaks and enabling timely responses to protect drinking and recreational water — especially for vulnerable groups such as children^34,35^ and pets^36–39^.

Other detection methods include fluorescence, spectroscopy, and biosensors^40–48^ such as biological nanopores^49,50^. Nanopore sensing allows single-molecule detection by monitoring ionic current fluctuations as molecules interact with a biological pore. Recent advances in nanopore research have shown that three of the most abundant MC congeners (MC-LR, -YR, and -RR) can be distinguished by their unique current signatures using the α-hemolysin (α-HL)^49,50^ nanopore. However, these measurements required relatively high concentrations (1 μM^50^, 200 nM^49^), well above the WHO safety threshold.

Here, we present a nanopore-based approach for the simultaneous detection of MCs at concentrations below the second strictest WHO threshold (<12 nM, **Table S1**) by using wild-type aerolysin (WT AeL) – a pore-forming toxin known for its ability to distinguish (bio)polymers, sugars, peptides with single amino acid differences, and various post-translational modifications^51–55^. We achieved picomolar detection of MC-LR at levels well below the strictest WHO threshold (<1 nM) by using asymmetric salt gradient and showed the capability of discriminating MC congeners while in mixture in a lake water background at an environmentally relevant concentration. Moreover, combining experiments and molecular dynamics simulations, we identify how single-point mutations influence the sensitivity of aerolysin, shedding light on its mechanism for the microcystin discrimination. Our results set the basis to use biological nanopores for real-time detection and monitoring of the presence and abundance of microcystins in rivers and lakes.

### Characterization of MC-LR, MC-RR, MC-YR with wild-type aerolysin nanopore

Single-channel recordings using WT AeL nanopore were conducted to investigate current blockage characteristics and assess the translocation of MCs. The WT AeL transmembrane barrel is formed by the oligomerization of seven identical protomers into a heptameric pore^56–58^. The experimental setup (**Fig. 1B**) comprised *cis* and *trans* electrolyte-filled chambers, separated by a membrane and connected by a single nanopore. The key constriction sites, R220 and K238, are known to be critical for sensing applications^51,52,54,57,59–61^. Upon applying an electrical bias, a stable ionic current (open pore current, *I₀*) is generated. Introduction of an analyte into the *cis* chamber induces transient current blockades, characterized by their magnitude (*I*), dwell time (the duration the analyte spends in the pore), and event frequency — parameters that provide insights into the analyte’s molecular structure and translocation dynamics.

The selected MCs exhibit structural variability at positions 2 (X) and 4 (Z), as shown in **Fig. 1A**. The 3 most studied congeners of MCs differ by a single residue at the X site, where either a leucine, arginine or tyrosine (L, R, Y) residue can be found. Initially, we focused on congeners that solely differ at the X site, while maintaining a fixed arginine residue at site Z. These congeners, denoted as MC-LR, MC-RR, and MC-YR, exhibit net charges of -1, 0, and -1 at pH 7.5, respectively.

Experimental conditions were adapted from previous studies^50,62^, using 4 M KCl to enhance capture and translocation of neutral, negatively and positively charged peptides through WT AeL by increasing electro-osmotic flow (EOF) and reducing electrostatic interactions^62^ . This approach enabled sensitive detection of MCs at 50 nM — substantially lower concentrations than previously used in the α-HL nanopore studies^49,50^.

As illustrated in **Fig. 1C**, 50 nM concentrations of MCs yielded clear and prolonged blockade events (**Figs. S1–S3**). Current blockage and dwell time served as the primary parameters for congener differentiation. Mean dwell times were calculated from exponential fits of dwell time distributions. Scatterplots of dwell time vs. current blockage (**Figs. S5–S8**) were generated across voltages from 60 to 200 mV to determine translocation thresholds, inferred from decreasing dwell times with increasing voltage. A voltage of 100 mV provided optimal separation among the 3 congeners (**Fig. 1D**), with MC-RR showing 6.6% and 8.7% lower current blockage than -YR and -LR, respectively.

MC current blockage followed the trend: MC-RR < -YR < -LR, with MC-YR and -LR differing by only 2.1%. Dwell time analysis (**Fig. 1E**) showed that MC-LR and -YR, both negatively charged, began translocating at lower voltages compared to neutral - RR. The longest dwell time for MC-RR (314.2 ± 66.4 ms) occurred at 160 mV, indicating that higher voltage is needed to drive its translocation due to its neutral charge.

Although larger molecular volume is often associated with greater current blockage, no systematic correlation was found between the volume of the variable amino acid and the respective congener’s current blockage (**Table S2**). Despite arginine’s (R) larger volume (173.4 Å³) compared to leucine (L) (166.7 Å³)^63^, MC-RR generated less blockage than -LR, consistent with prior α-HL data. Similarly, MC-YR, with a larger tyrosine residue (Y) (193.6 Å³), showed less blockage than -LR. This suggests that factors beyond volume — such as charge, conformation and hydrophobicity — influence blockade behavior. MC-RR showed the smallest blockage, likely due to its +1 charge, compared to -LR and -YR, promoting intramolecular interactions and reducing the interaction with AeL (also reflected in **Fig. 1E** by short dwell times), likely resulting in a more compact conformation in the long and narrow β -barrel of AeL.

The increasing blockage from MC-RR to -YR and -LR aligns with their rising negative charge and hydrophobicity. RR, being neutral and least hydrophobic, interacts weakly with the pore, whereas LR, the most hydrophobic and negatively charged, exhibits stronger binding and longer dwell times. YR shows intermediate behavior consistent with its physicochemical profile.

NMR spectroscopy was used to probe the conformational states of the 3 MCs in solution by comparing their backbone ^1^H and ^13^C chemical shifts, which are exquisitely sensitive to conformation (**Table S3**). These data show that MC-LR and - YR must be structurally more similar to each other than to -RR (**Fig. S9**). In particular, the ^13^Cα and ^13^Cβ chemical shifts at E6 defer markedly between MC-LR and MC-RR (^13^C-HSQC spectra were not attainable for MC-YR due to its low concentrations, but ^1^H-TOCSY patterns are almost identical to MC-LR’s). Although explicit interpretation of the chemical shift changes is not trivial, the higher Δδ_Cα_ - Δδ_Cβ_ in MC-RR would indicate a slightly higher level of local structure. Molecular dynamics simulation for MCs in solution^64^ have shown that these are very flexible molecules, and it appears from this NMR data that the conformational landscape is altered in MC-RR compared to MC-LR and MC-YR. These differences would be consistent with the trends found in current blockages (**Fig. 1D**), and if it were at the root cause of the observed trends, it would imply overall somewhat tighter states in MC-RR. However, what was not observed in the NMR spectra — but becomes evident in the nanopore experiments between 140 mV and 200 mV (**Figs. 3A, S6, S7**) — is the presence of two distinct populations for MC-RR, suggesting that the congener may adopt multiple orientations and conformations upon entering and interacting with the pore, as similarly observed in MD simulations of MCs in solution^64^.

Finally, a mixture experiment tested whether WT AeL could resolve all three congeners simultaneously. An initial binary mixture of MC-RR and -YR (**Fig. 1F**) was followed by -LR addition (**Fig. 1G**) at adjusted concentrations (25 nM MC-LR, 50 nM - RR, 20 nM -YR) to balance capture rates to ensure uniform signal distribution (**Fig. S4**). The resulting current blockage distributions aligned well with the individual experiments, confirming that WT AeL can distinguish MC congeners differing by a single residue.

The capture frequency of the negatively charged congeners MC-LR and -YR increased more rapidly with increasing voltage, compared to neutral -RR (**Fig. S4**). This might be due to the combined effect of electrophoretic mobility and electroosmotic flow (EOF) leading to an increased frequency of negatively charged species at higher voltages, whereas neutral species are less affected by electrophoretic force^60,65^.

## Dissecting the microcystin translocation mechanism

To gain more insights into which region of the pore influences the current blockage the most, we compared results from nanopore experiments and molecular dynamics (MD) simulations (**Fig. 2A**), by measuring current blockages (in experiments and simulations) and dwell times (from experiments only) for MC-LR (and MC-RR, -YR in **Fig. S10**) in WT AeL and comparing with two mutants that alleviate the two constriction points, K238A and R220A (**Fig. 2B and 2C**). In the simulations, the MC-LR molecule was placed inside the β-barrel, in-between the two constriction points, near K238 – a conformation we hypothesized is dominant for producing MC-LR current blockages (**Fig. 2A**) – spanning residues from S228/Q268 to G234/N262 (residue numbers correspond to the two β-strands of AeL’s barrel) and with no MC atoms within 4 Å of any protein atoms. In both experiments and simulations, currents were measured in symmetric 4 M KCl buffer.

**Figure 2.**
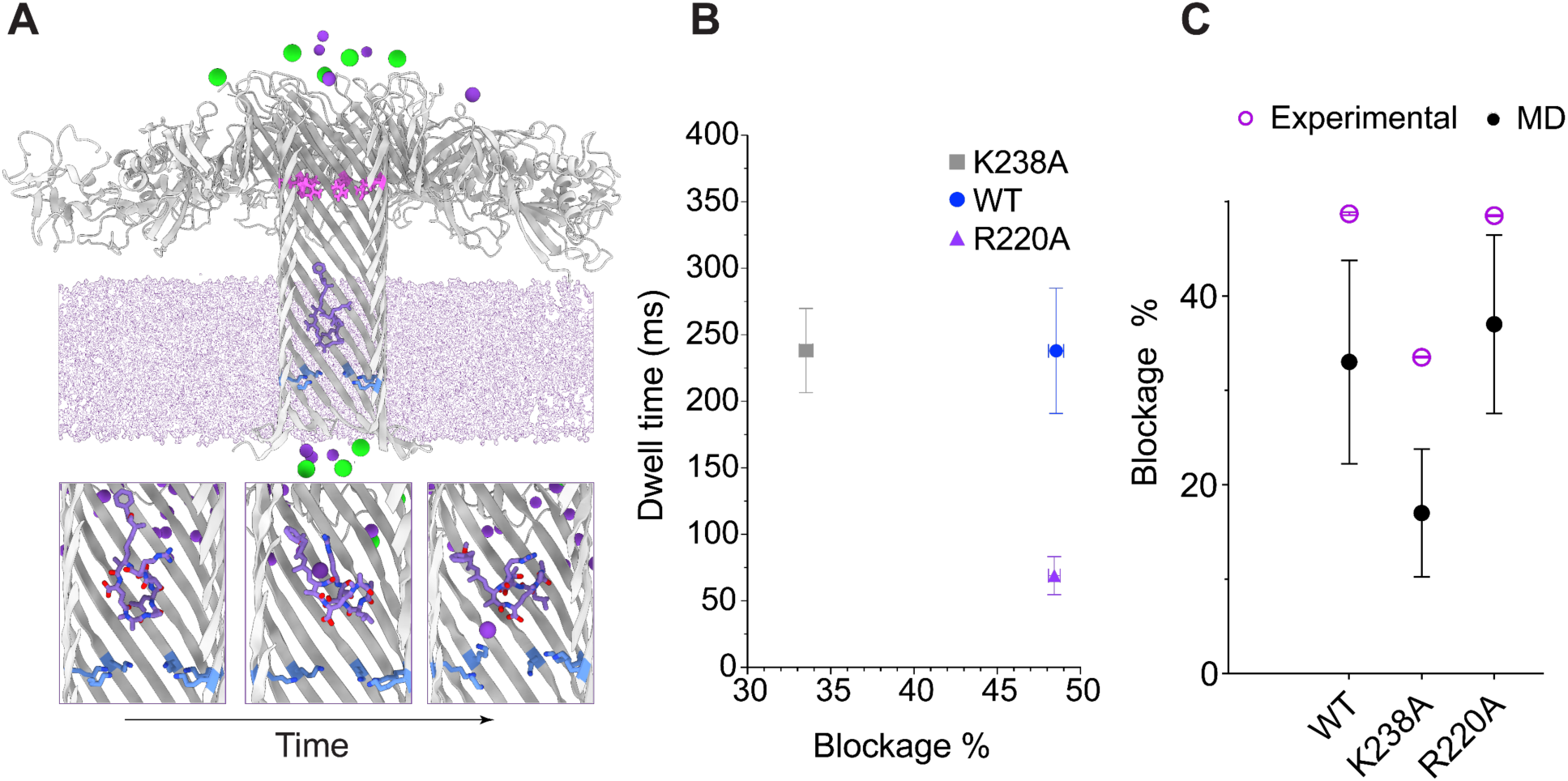
Dissecting MC-LR blockage with mutagenesis and MD simulations. **(A)** MD setup (top) and changes in MC-LR orientation within the pore over time (bottom). Cartoon of WT AeL with highlighted R220 (magenta), K238 (blue) constrictions; MC-LR structure (purple); K^+^, Cl^-^ ions (dark purple, green spheres, respectively) are visualized. Cartoon of WT AeL showing R220 (magenta) and K238 (blue) constrictions, MC-LR (purple) placed inside the β-barrel, and visualized K⁺ (dark purple) and Cl⁻ (green) ions. As demonstrated by the frames of MC-LR from the beginning and end of the simulation run, the orientation and conformation of -LR clearly changes. **(B)** MC-LR experimental dwell time and blockage comparison between WT AeL (48.5 ± 0.4 %), K238A (33.5 ± 0.4%) and R220A (48.4 ± 0.3%) mutants at 160 mV (at least 3 pores for each mutant were analysed). Experimental data acquired in 4 M KCl, buffered with 10 mM Tris, 1.0 mM EDTA at pH 7.5. **(C)** MC-LR experimental and MD predicted blockage comparison between AeL mutants at 160 mV.

**Figure 3.**
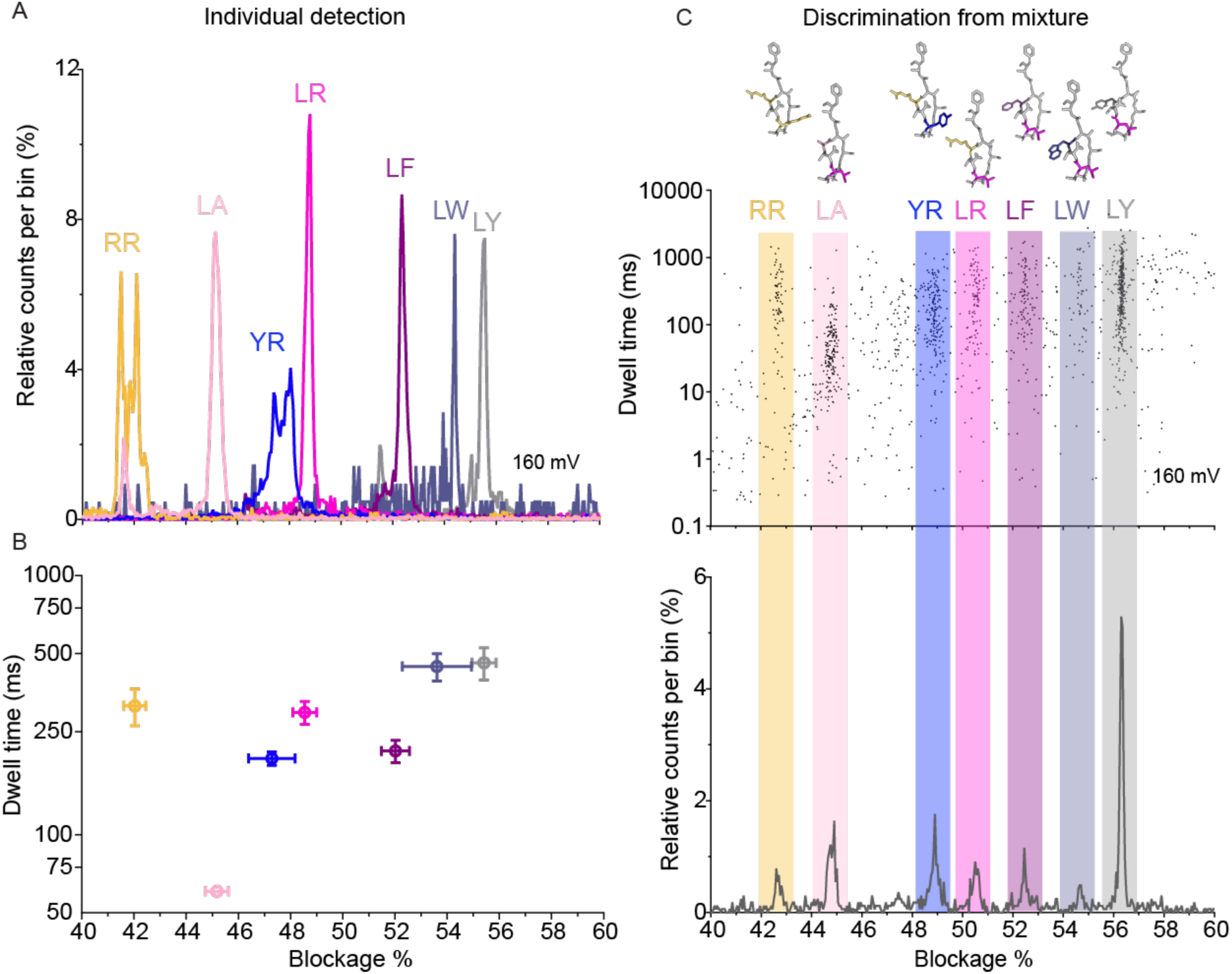
Expanding detection of MCs with WT Ael. **(A)** Each MC congener produces a distinguishable current blockage in individual experiments at nanomolar concentration at 160 mV represented in a histogram (n>500 events). **(B)** Discrimination of 7 congeners at nanomolar concentrations in individual experiments, based on current blockage and dwell time at 160 mV. **(C)** Representation of 7 MCs, a scatterplot derived from a single nanopore recording of a mixture containing the 7 MCs (top), and a resulting histogram (bottom) are shown. Data acquired in 4 M KCl, buffered with 10 mM Tris, 1.0 mM EDTA at pH 7.5.

Experimentally, MC-LR induced comparable levels of blockage in both WT and R220A pores, although the dwell times differed between the two pores (**Fig. 2B**). In contrast, the K238A mutant exhibited significantly reduced blockage compared to WT and R220A, but a similar dwell time as the WT AeL pore. Based on these findings, the main current blockage (sensitivity) seems modulated only by the K238 constriction, as mutation of the R220 constriction does not alter the current compared to WT. This is further supported by MD simulations, which show a similar trend in blockage percentages across the three different pores (WT, K238A and R220A, **Fig. 2C**) when MC-LR was positioned within the pore cavity as stated above. In contrast, dwell time seems to be primarily regulated by the R220 constriction, since the R220A mutant exhibits a shorter dwell time for MC-LR while maintaining the same current blockage as WT.

## Expanding the detection range of microcystins

To further test the ability of WT AeL to discriminate MCs, the electrical readouts of four additional MC congeners were recorded. These congeners possess a fixed leucine residue at position X (**Fig. 1B**) and a variable residue at position Z, being alanine, phenylalanine, tyrosine or tryptophan, depicted in **Fig. 3** as MC-LA, -LF, -LY and -LW respectively. Under a voltage of 160 mV, we were able to separate all the tested MC congeners, as shown by the current blockage histograms obtained from individual experiments (**Fig. 3A, S12**). More hydrophobic MCs, especially MC-LF and -LW, produced significantly longer dwell times at 160 mV compared to other MCs (**Fig. S11**). The dynamic range of our method is reliably achievable up to 50 nM for the tested microcystins, except for MC-LY and -LW, for which optimal signal can only be observed at concentrations up to 25 nM. This range is suitable for detecting environmentally relevant concentrations, as MC levels during cyanobacterial blooms rarely exceed 50 nM^13,66–72^. MC-LY and -LW required testing at 25 nM because, at higher concentrations (e.g., 50 nM), they frequently produced two-level blockade events (**Fig. S13**) that hindered rapid and confident identification. These events often failed to return to the open-pore current, particularly at lower voltages (e.g. 100 mV). A similar behavior was initially observed for MC-LR at 2 μM (**Fig. S14**), which was resolved by lowering the analyte concentration. Based on these observations, we propose two hypotheses to explain the two-level events observed with MC-LY and -LW. First, they may result from strong aromatic interactions (e.g., π–π or cation–π) between the MCs and the pore^73^, resulting in MC getting stuck in 2 possible states within the pore. Secondly, two MC-LY or -LW (and similarly -LR at 2 μM) molecules may be entering the pore sequentially or in a chain-like manner, potentially driven by intermolecular aromatic interactions that increase affinity and facilitate cooperative entry and exit dynamics.

Bulkier MCs such as MC-LF, -LY, -LW provided deeper pore blockage than -LA, seemingly the smallest MC based on structure. However, MC-LA produced deeper blockage than -RR (**Fig. 3A**), probably due to its higher hydrophobicity compared to RR as previously discussed. Separation of the 7 MC species was clearly evident in dwell time and current blockage dimensions, without the need for further sophisticated analysis (**Fig. 3B**). We observed that all the MCs did possess a similar dwell time for this voltage except MC-LA, probably due to its smaller size. The results demonstrate that the main parameter to discriminate MC is current blockage, since very narrow current distribution can be observed. To go a step beyond, a mixture experiment with all 7 MC congeners present in a single sample was performed, relevant for real-world applications (**Fig. 3C**). As shown, the individual current slightly shifted, but the specific population corresponding to each MC remained conserved and did not overlap, demonstrating the capabilities of WT AeL to discriminate up to seven MC congeners simultaneously.

Conserved leucine is indicated in pink, and the variable amino acid is colored according to the histogram. Molecular properties of MCs are shown in **Table S2.**

## Discrimination of microcystins in lake water

Discrimination of up to 7 MCs was achieved in lab-controlled conditions; subsequently, we focused on assessing MC detection in natural waters such as lakes or rivers. For that, we assessed the detection of MC-LR and -YR in water sourced from Lake Geneva, Switzerland. This condition was anticipated to pose a challenge due to the minimal difference in current blockage (2.1% from **Fig. 1D**). The lake water, confirmed to be free of MCs (as determined by ELISA; see **Methods**), was used as a background matrix. MC-LR and MC-YR were spiked at 25 nM and 20 nM, respectively – concentrations resembling those found in lake environments affected by strong cyanobacterial blooms^45^. Lake water background signals were recorded in unspiked samples. As shown in the scatterplots in **Fig. 4A, E, I**, lake water did not produce any significant background and mainly short events could be observed. More importantly, there were no events within the current blockage windows expected for MCs, which could otherwise interfere with MC discrimination.

**Figure 4.**
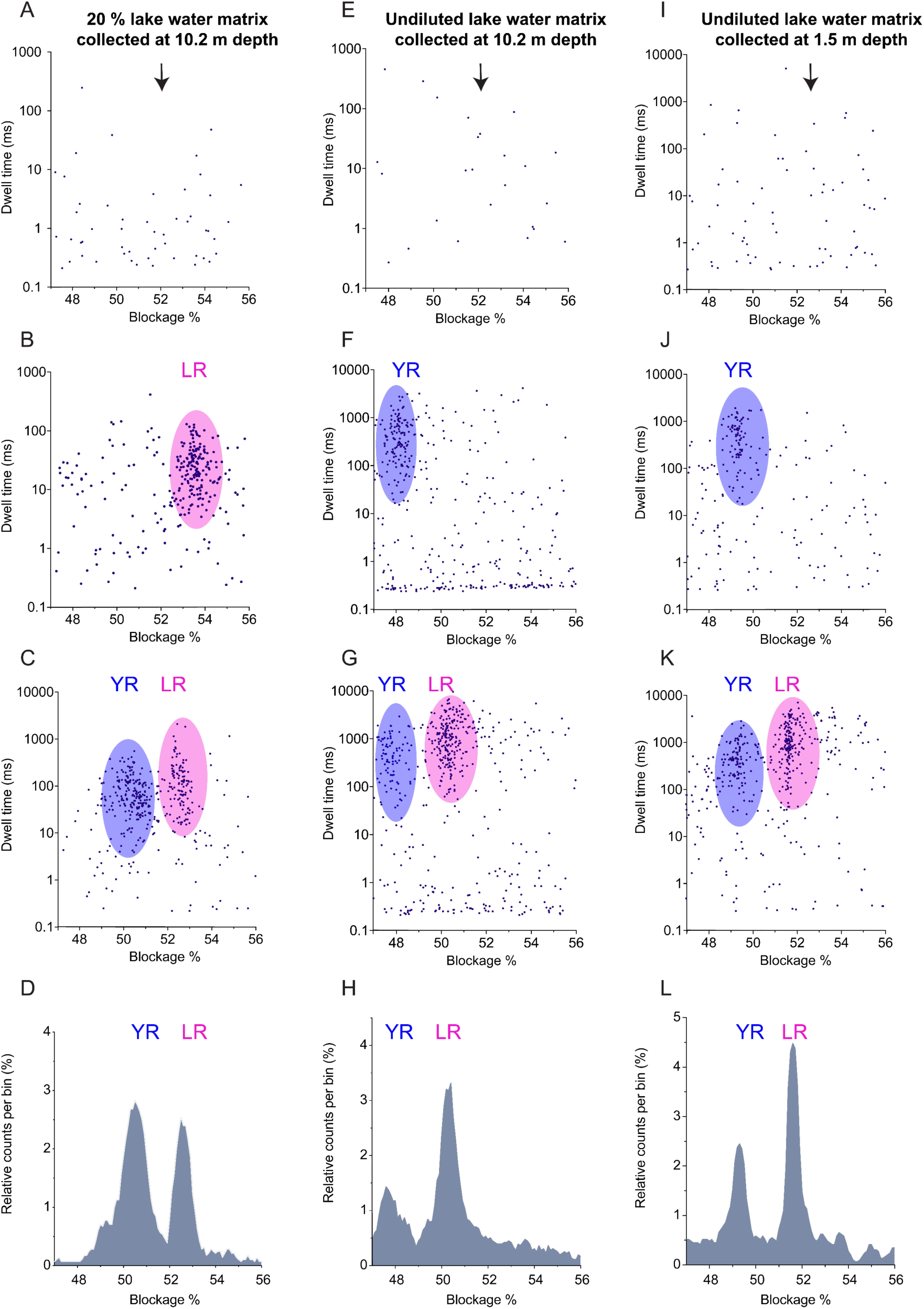
Discrimination of MCs in a lake water matrix. (A-D) Top to bottom: Diluted matrix, containing 20 % lake water collected at 10.2 m depth, the scatterplots of the single pore upon sequential addition of MCs to the matrix, histogram (bin size = 0.1) of the mixture experiment in C, respectively. **(E-H)** Undiluted lake water background collected at 10.2 m depth, the scatterplots of the single pore upon sequential addition of MCs to the matrix, histogram (bin size = 0.1) of the mixture experiment in G, respectively. **(I-L)** Undiluted lake water background collected at 1.5 m depth, the scatterplots of the single pore upon sequential addition of MCs to the matrix, histogram (bin size = 0.1) of the mixture experiment in K, respectively. Data shown recorded at 100 mV, 3.3 M KCl for **A-D**, and 4 M KCl for **E-L**.

First, MC-LR, and -YR were spiked into a diluted lake water matrix (at 3.3 M KCl instead of the 4M KCl previously used, see **Methods** for water sample preparation) and the same resolution in current blockage was retained as observed in **Fig. 1G**, with a separation of approximately 2.5% between MC-YR and -LR (**Fig. 4A-D**). Next, we directly used water collected at 10.2 m (**Fig. 4E**) and 1.5 m depths (**Fig. 4I**), representing water from the deep chlorophyll maximum layer and the surface layer, respectively. To keep similar experimental conditions and assess the background noise resulting from any other (in)organic material present, the same electrolyte concentration (4 M KCl) was kept for the following experiments, and the MC-YR, -LR were spiked sequentially (**Figs. 4F-H, 4J-L**). Assessing this under high-salt conditions is necessary, as elevated salt levels can enhance the capture of any molecules from lake water, potentially contributing to signals that interfere with MC detection. Sequential spiking of the MCs to the respective samples demonstrated that the current separation was preserved (**Fig. 4H, L)**. As shown in **Fig. 4D, H**, **L**, the current separation was still present as in individual and mixture experiments performed previously (**Fig. 1D**). A slight shift in current is observed, which can be attributed to the different ion composition present in the lake water, but with no apparent consequence for the separation of MC-LR and -YR (2.4 ± 0.2 % compared to 2.1 % in **Fig. 1D**). These results demonstrate that AeL nanopore enable reliable and selective detection of these microcystin congeners in natural water samples, at environmentally relevant concentrations during blooms, without interference from the sample matrix.

## Pushing the microcystin detection limit to reach picomolar concentrations

To further enhance the sensitivity of the nanopore-based detection system, a series of experiments was performed to determine the lowest achievable limit of detection (LOD) and quantification for MC-LR^31^. Our aim is to detect concentrations well below the WHO’s lowest guideline value of 1 nM for chronic lifetime exposure in drinking water^74^, as well as environmentally relevant concentrations typically found in lake surface waters during bloom events ^10^. In concentrated cyanobacterial scums, MC levels can even reach micromolar concentrations (up to 14000 µg/L of total MC)^75^. However, in open surface or lake waters, MCs are usually reported in the nM range or lower (ng/L to low µg/L). For example, microcystin levels can peak between 105 pM to 147 pM^13^ in Lake Geneva to approximately 1 nM (1,000 ng/L) in other Swiss lakes^66,67^, 10 nM in highly eutrophic lakes in the USA^68^, 24 nM in the southern US Reservoir^69^, 46 nM in Hoan Kiem Lake (Vietnam)^70^, 50 nM in Lake Taihu (China)^71^, and up to 78 nM in lakes in Mexico^72^. To improve the detection, we employed a salt concentration gradient, a technique previously demonstrated to increase event frequency for ssDNA^51,76^. Specifically, we used asymmetric salt condition with 0.15 M KCl buffer solution in the *cis* chamber and 4.0 M KCl buffer solution in the *trans* chamber (**Fig. 5**).

**Figure 5.**
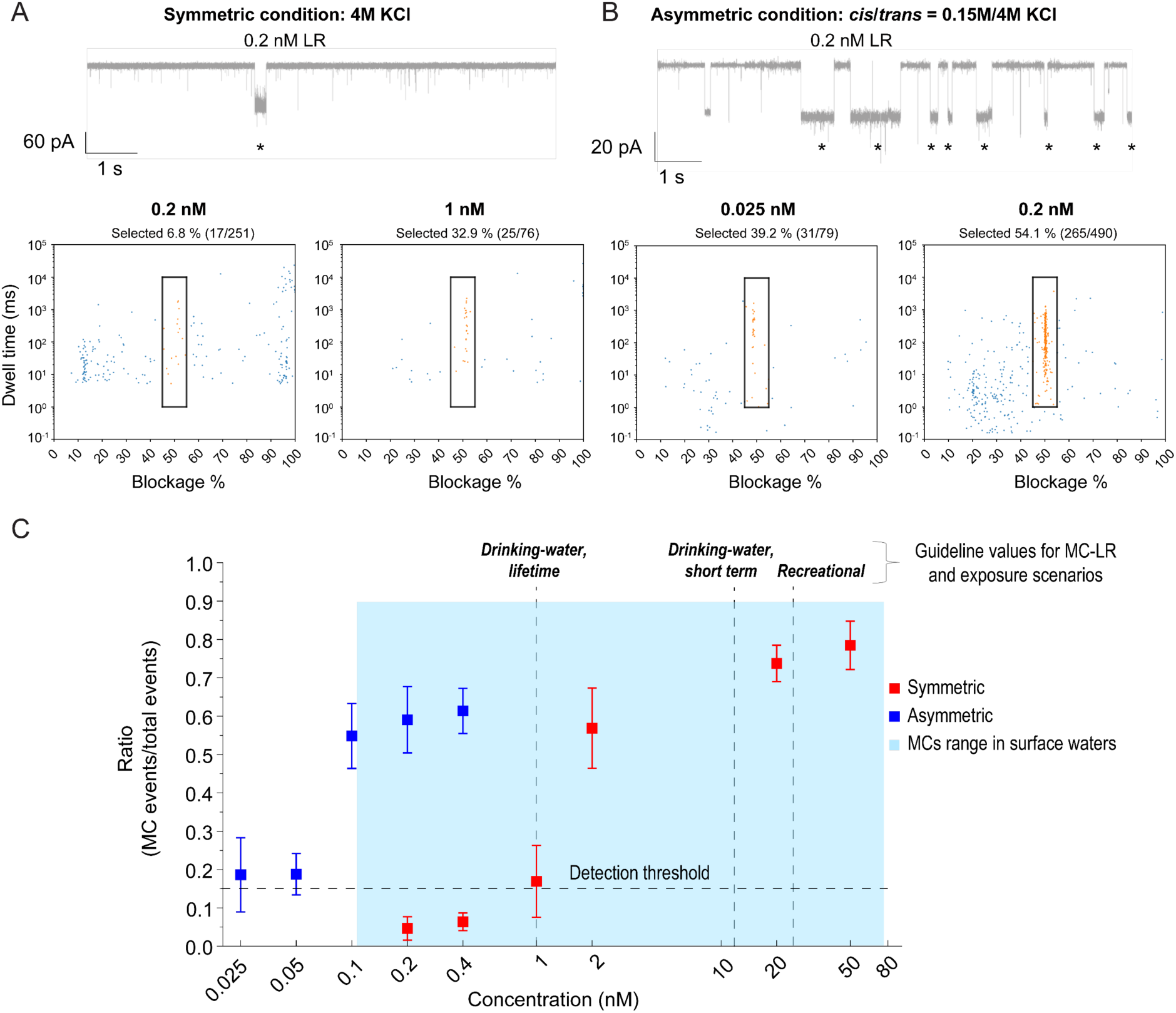
Probing the microcystin detection limits. **(A)** Raw current trace, where MC-LR events are marked with a star, and scatterplots in symmetric conditions at 0.2 nM and 1 nM MC-LR. **(B)** Raw current trace, where MC-LR events are marked with a star, and scatterplots in asymmetric condition at 0.025 nM and 0.2 nM MC-LR. **(C)** Ratio of MC-LR events selected from scatterplots towards background events in symmetric (blue) and asymmetric conditions (red). Ratio of 0 shows that only background events are detected whereas a ratio value close to 1 indicate that most events detected are from MC-LR. Data acquired at 100 mV in their respective KCl concentrations, buffered with 10 mM Tris, 1.0 mM EDTA at pH 7.5. Shaded area (light blue) represents the typical range of MC concentrations observed in surface waters^13, 59–72^. The three WHO guideline values for MC-LR, corresponding to different exposure scenarios, are indicated above.

**Figure 5** shows two representative raw current traces for 200 pM MC-LR in symmetric (**A**) and asymmetric conditions (**B**). The number of events (depicted by a star) is drastically improved in asymmetric conditions. This is further demonstrated in the scatterplots, where MC-LR events (highlighted within a dark rectangle) account for 54.1% of total events under asymmetric conditions, compared to only 6.8% under symmetric conditions. The LOD was defined as the lowest concentration at which the event ratio exceeded the detection threshold, set at the ratio of 0.15, i.e. three times from the ratio observed at the analyte concentration, where MC-LR was barely detectable (0.2 nM at symmetric salt condition, Fig. 5A).

Additionally, event frequency (in Hz) was calculated by fitting inter-event time in **Fig. S15**. Therefore, concentrations of MC-LR ranging from 25 pM to 50 nM were screened under both symmetric and asymmetric conditions, and the ratio of events corresponding to microcystin was extracted (from 45 % - 55 % current blockage; from 1 ms - 10 s range; Fig. 5A, B; **Table S4**) and compared to all the events detected. Based on the scatterplots (Fig. 5A) and the ratio (Fig. 5C), under symmetric conditions, MC-LR was confidently detected at concentrations as low as 1 nM, accounting for 32.9 % of observed events. In contrast, a comparable event ratio (39.2%) was still observed at 25 pM under asymmetric conditions (Fig. 5B), representing a 40-fold improvement in detection sensitivity. The event ratio at 2 nM under symmetric conditions was similar to that observed at 100 pM under asymmetric conditions, representing a 20-fold improvement in sensitivity. Using this approach, we can therefore significantly improve the detection capability of WT AeL, enabling reliable MC-LR detection at concentrations substantially lower than the 1 nM WHO guideline with an LOD of 25 pM for asymmetric condition and 1 nM for symmetric condition. This sensitivity is well below the concentrations for water safety and environmental monitoring, demonstrating the strong potential of this method for real-world applications.

## Conclusion

In conclusion, using the aerolysin nanopore, we demonstrate real-time detection of 7 MC congeners with high sensitivity and specificity competing with state-of-the-art LC-MS/MS technique. We successfully detected the congeners at nanomolar concentrations and achieved picomolar detection for MC-LR under asymmetric buffer conditions. This represents a significant advancement over previous studies and positions nanopore-based detection as an effective technology for real-world applications in water surveillance. The method’s ability to discriminate seven MC congeners within mixture experiments, including those spiked in lake water, underscores the reliability and robustness of the aerolysin nanopore platform for direct, real-time monitoring of microcystins.

Beyond microcystins, water sources worldwide are threatened by a diverse array of contaminants, including other cyanotoxins (such as anabaenopeptins, cylindrospermopsins, anatoxins or saxitoxins), pesticides, pharmaceuticals, heavy metals or other industrial pollutants^14,15,77^. The present study highlights that the aerolysin nanopore can be in principle used as an analytical tool for the detection of these water contaminants. Through protein engineering or chemical modifications, aerolysin’s sensitivity and selectivity can be further expanded to cover sensing of a wider range of (cyano)toxins or small molecules, as shown in other domains for nanopores such as MspA or α-hemolysin^78^. Moreover, the adaptability and compact nature of the nanopore technology makes it ideal for integration into portable, field-deployable devices for on-site water quality monitoring^79–81^.

Lastly, the ability to rapidly and accurately monitor multiple toxins in real time is crucial for safeguarding water quality and protecting public health. Our results pave the way for more comprehensive and responsive water quality management strategies, addressing critical environmental and public health concerns associated with cyanotoxins and other pollutant contamination.

## Authors contributions

A.A., T.K. and M.D.P. conceived the project. A.A., J.F.B.J., T.K. and M.D.P wrote the original draft, all authors contributed to the final manuscript; A.A. and J.F.B.J. performed and analyzed nanopore experiments; M.J.M. produced aerolysin pores; L.A.A. performed and analyzed NMR experiments and MD simulations; C.C. provided guidance in the analysis of the nanopore data; A.C. collected the lake water samples, E.M.L.J. advised on handling of microcystins; A.C. and E.M.L.J. provided insights and contributed to the interpretation of the environmental aspects of the project.; T.K. and M.D.P. supervised the project and acquired funding.

## Acknowledgments.

We thank the Central Environmental Laboratory at the Institute of Environmental Engineering (EPFL, CH) and in particular Dr. F. Breider and D. Grandjean, for their support in ensuring the quality of the tested and commercially sourced microcystin congeners by LC/MS-MS. We thank S. Vacle for contributing to the production of aerolysin pores. We acknowledge the contributions of A. Bayramova, J. S. Behler, and M. J. Morella to the single-channel recording experiments. We thank LéXPLORE platform for access and support that enabled the collection of water samples. We thank Dr. V. Rougé from the laboratory of PD Dr. E. M. L. Janssen (EAWAG, CH) for fruitful discussions on the handling of microcystins and their analysis. We thank the laboratory of Prof. A. Radenovic (EPFL, CH) for providing the equipment used in the asymmetric salt experiments, and M. F. Mitsioni and Dr. S. F. Mayer for their assistance with the experimental setup and manipulation. This work was supported by an EPFL iPhD grant to M.D.P. and T.K. M.D.P. acknowledges the Swiss National Science Foundation (SNSF, grant number 200021L_212128); for C.C the SNSF (PR00P3_193090). LAA and MDP acknowledge the Swiss Supercomputing Center CSCS for HPC time.

## Methods

### Preparation of aerolysin

The wild-type aerolysin sequence was cloned into the pET22b vector with a C-terminal hexahistidine tag as in previous study ^82^, followed by expression and purification ^61^ from *Escherichia coli* BL21 DE2 pLys cells. The bacterial culture was cultivated in Luria-Bertani medium until it reached an optical density between 0.6 and 0.7. Protein expression was then induced with 1 mM isopropyl β-d-1-thiogalactopyranoside (IPTG), followed by overnight incubation at 20 °C. The harvested cells were suspended in a buffer containing 20 mM sodium phosphate (pH 7.4) and 500 mM NaCl and mixed with cOmplete Protease Inhibitor Cocktail (Roche). Cells were lysed by sonication, after which the lysate was centrifuged at 20,000xg for 35 minutes at 4 °C. The clarified supernatant was loaded onto a pre-equilibrated HisTrap HP column (GE Healthcare) in the same buffer. Protein elution was performed using a 40-column volume gradient of buffer containing 20 mM sodium phosphate (pH 7.4), 500 mM NaCl, and 500 mM imidazole. Subsequently, the eluted protein underwent buffer exchange into a final solution of 20 mM Tris and 500 mM NaCl (pH 7.4) using a HiPrep Desalting column (GE Healthcare). The purified protein was then rapidly frozen in liquid nitrogen and preserved at -20°C for future use.

### Nanopore single-channel measurements

Powdered 1,2-Diphytanoyl-*sn*-glycero-3-phosphocholine (DPhPC) lipid (Avanti Polar Lipids, Inc., Alabaster, AL, USA) was dissolved in octane (Sigma-Aldrich Chemie GmbH, Buchs, Switzerland), to get a final concentration of 8 mg/mL. The purified protein was diluted to a concentration of 0.2 μg/mL and activated using Trypsin-agarose (Sigma-Aldrich Chemie GmbH, Buchs, SG Switzerland) for 2 hours at 4 °C. Subsequently, the solution was centrifuged (11,000 rpm, 5 min, 4 °C) to remove trypsin.

Single-channel current recording experiments were performed on an Orbit Mini workstation (Nanion, Munich, Germany), using MECA 4 recording chambers with an 50 µm cavity size. DPhPC membranes were formed across the four cavities of the chamber, separating the well into *cis* and *trans* compartments. Each microcavity contains an integrated Ag/AgCl microelectrode through which an electrical bias is applied. The heptameric pore is formed upon addition of aerolysin to the *cis* chamber, followed by its self-assembly and insertion into the membrane. The experiments were performed at 25 °C under symmetrical buffer condition with both compartments filled with a 150 µL electrolyte solution, where MC analytes were added to the *cis* compartment. All data was gathered using a sampling rate of 100 kHz and filtered with a 5 kHz low-pass filter. In order to obtain relatively uniform capture between MCs, the ratio of MC-LR : -RR : -YR used for the mixture was chosen based on the capture rate extracted from individual MC experiments (**Fig. S4**). Since the ratio of MCs expected to result in similar capture was varying depending on voltage, that of 160 mV was chosen.

To perform asymmetric salt experiments, Teflon films with 50 μm diameter openings were secured between two compartments of a Teflon chamber using high-vacuum grease (Dow Corning Corporation, Midland, MI, USA). *Cis* and *trans* compartments were connected solely through the aperture in the film, with each side containing an Ag/AgCl electrode. Both sides of each aperture were pretreated with 1 μL of a 1 % (v/v) hexadecane (Sigma-Aldrich Chemie GmbH, Buchs, Switzerland) solution in hexane (Sigma-Aldrich Chemie GmbH, Buchs, Switzerland), after which the chamber was placed in the recording apparatus. DPhPC bilayers were then established across the aperture using the folding technique described in earlier protocols^83,84^. Electrolyte solution was introduced into both chambers, ensuring that the liquid remained below the level of the aperture. Lipids, dissolved at 10 mg/mL in pentane (Sigma-Aldrich Chemie GmbH, Buchs, Switzerland) were then applied to the electrolyte surface in each compartment. Once the pentane evaporated, the electrolyte volume was raised above the aperture to facilitate bilayer formation. The integrity of the resulting lipid bilayer was assessed by measuring its capacitance. When an analyte was added, the *cis* compartment was mixed by pipetting to ensure thorough distribution. Electrical currents were recorded at a sampling rate and a low-pass filter of 100 kHz using an Axopatch 200B amplifier (Molecular Devices, LLC., San Jose, CA, USA).

### Current signal processing

Information regarding open pore current (*I_0_*), mean residual current (*I*), average relative current (*I/I_0_*), event dwell time (*dwt*) and inter-event time - were extracted from the raw current traces and processed with in-house developed Python-based pipeline^61^. Current blockage was calculated as the difference between the open pore current (*I_0_*) and the current produced by the passage of the analyte through the pore (*I*), expressed as *ΔI/I_0_* (where *ΔI = I_0_-I*), and reported as ‘Blockage %’ in the figures. The raw signals are divided according to voltage discontinuities and significant time-scale shifts to identify stable single pore readings and discard segments where the pore self-blocks or where several pores are present. For each identified segment, the distribution of the open pore current is analyzed by applying a Gaussian fit to the peak of the current distribution that exhibits the highest mean value. Event detection is performed using a dual-threshold approach based on the open-pore current distribution: 1) for event initiation, a threshold of 7 σ below the mean open-pore current is applied, where σ represents the standard deviation of the open-pore current, 2) for event termination, a threshold of 3 σ below the mean open-pore current is applied. The average relative current is translated into the pore blockage % (ΔI/I_0_), the processed data is initially visualized as scatterplots, enabling the selection of events of interest (falling in the blockage range of 40 %-60 %, dwt range of 0.2 ms – 10 s) for further in-depth analysis, where both, current and dwell time distributions, are identified by fitting a Gaussian and an Exponential function, respectively. MCs produced rather defined current blockage distributions, with a more spread dwell time distributions, the event selection windows of the respective MCs are shown in **Figs. S4-S8, S12**. For event frequency analysis of both ratio (event selection %) and inter-event time (Hz), selection was modified (to a 45% - 55 % current blockage and 1 ms - 10 s dwell time range). Due to the possible occurrence of two-level events, the frequency calculated from inter-event times (Hz) might not accurately reflect the true event rate at concentrations > 10 nM, as multiple events can overlap and be counted as a single inter-event interval, causing the measured frequency to plateau or even decrease at higher concentrations. Therefore, we relied on the ratio of observed events rather than inter-event time-derived frequency, which provided more consistent and interpretable results across conditions, resulting in a sigmoidal relationship between ratio and concentration. We observed an increase in event frequency with concentration up to 2 nM; however, data above 10 nM were excluded from analysis, as the apparent decrease in frequency at higher concentrations was inconsistent with expected linear increase and likely reflected measurement artifacts (i.e two-levels).

### Lake water collection and sample preparation

Lake water samples were collected in early June 2022 from L’éXPLORE^85^ experimental platform of Lake Geneva (Switzerland). Samples were collected at the depths of 1.5 m and 10.2 m below the surface. Prior to nanopore experiments, all samples were filtered through a 0.2 μm membrane to remove particles, intact bacteria and other microorganisms. To validate the suitability of the matrices as negative controls for microcystin detection, an Enzyme-Linked Immunosorbent Assay (Biorbyt, UK) was performed to confirm the absence of microcystins. The diluted lake water matrix (20% v/v) shown in Fig. 4A**-D** was prepared by adding 30 μL of lake water to 120 μL of 4 M KCl, resulting in a final KCl concentration of 3.3 M.

### NMR spectroscopy

All NMR experiments were carried out in a Bruker 800 MHz spectrometer equipped with an Avance Neo console and a triple resonance cryoprobe, at 298 K. Samples were prepared in 20 mM phosphate (pH 5.9) buffer^86^ at concentrations close to 1 mM (MC-LR and MC-RR) or 0.1 mM (MC-YR). Resonances were assigned by combining ^1^H,^1^H TOCSY, ^1^H,^1^H NOESY, and ^1^H,^13^C HSQC experiments (the latter only for MC-LR and MC-RR), guided by assignments available in the literature for MC-LR^86–88^.

### MD simulations

We set up the systems for MD simulations by using our recently published high-resolution cryo-EM structure of WT AeL, noting in particular several key protonation states critical for proper simulation as described in the paper reporting said structure along with MD simulations^57^. For this work we parametrized the systems and carried out the simulations by using the same CHARMM-GUI-based protocol, utilizing the same DPhPC membrane but with a solution of 4 M KCl as employed in the experiments. The MC molecules were parametrized with the CHARMM General Force Field^89^. The simulations were run with Gromacs 2024 by using the standard production stage scripts produced by CHARMM-GUI (pressure coupling to 1 atm, a temperature of 303 K, 2 fs integration steps, LINCS-based restraints on hydrogens, and PME electrostatics with 12 Å cutoff), but applying the indicated voltages along the z dimension, after the standard minimization and equilibration protocol obtained from CHARMM-GUI. The simulations were visualized with VMD and processed with custom scripts to obtain currents by counting the numbers of K^+^ and Cl^-^ ions passing per unit time, from which the blockages were then calculated.

**Supplementary information**. Supplementary materials, **Supplementary Tables S1-S5** and **Figs. S1-S15**.

