## Supplementary information for "Sub-Nanomolar Detection and Discrimination of Microcystin Congeners Using Aerolysin Nanopores"

### Materials

All reagents used were of analytical-grade quality. The stock solutions of microcystins (Enzo Life Sciences, Farmingdale, NY, USA) at 1 mM (MC-LR, MC-RR), 200 μM (MC-YR, MC-LA, MC-LF, MC-LY, MC-LW) were dissolved in ultrapure water, and stored at −20 °C. Electrolyte solution was composed of 4M KCl (Sigma-Aldrich Chemie GmbH, Buchs, Switzerland), 10 mM Tris (Fischer, UK), 1.0 mM EDTA (Fisher, UK), and the pH was adjusted to 7.5. Prior to single-channel experiments, stock solutions of MCs at 200 nM in the respective electrolyte buffer were prepared.

### Supplementary Figures

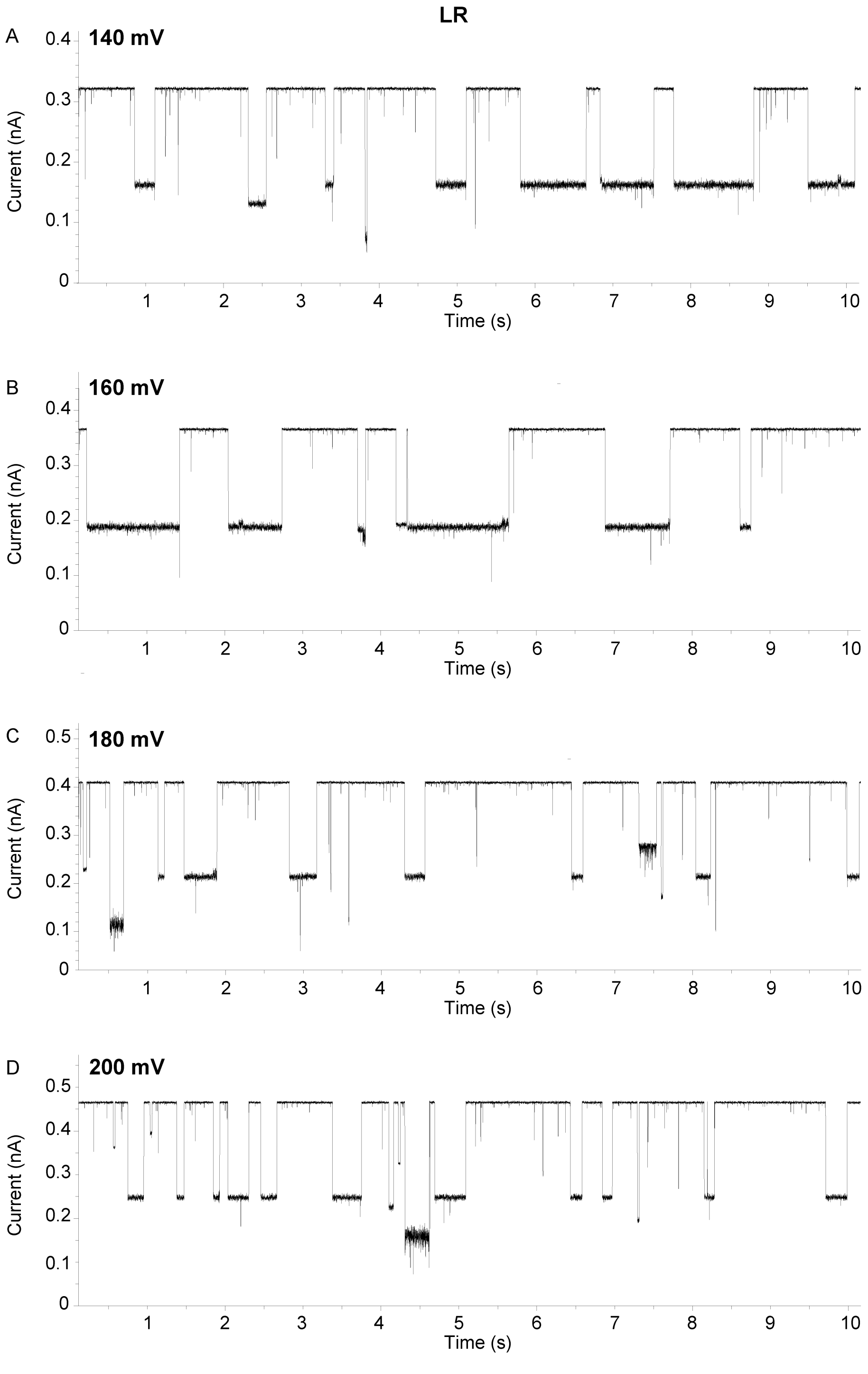


**Figure S1. Raw current traces of MC-LR.** Raw current traces at 140mV **(A)**, 160 mV **(B)**, 180 mV **(C)**, 200 mV **(D)** acquired in 4 M KCl, buffered with 10 mM Tris, 1.0 mM EDTA at pH 7.5; traces were filtered to 500 Hz. Increasing voltage results in shorter dwell times, as shown in the traces. Data extraction was performed as described in the Methods section.

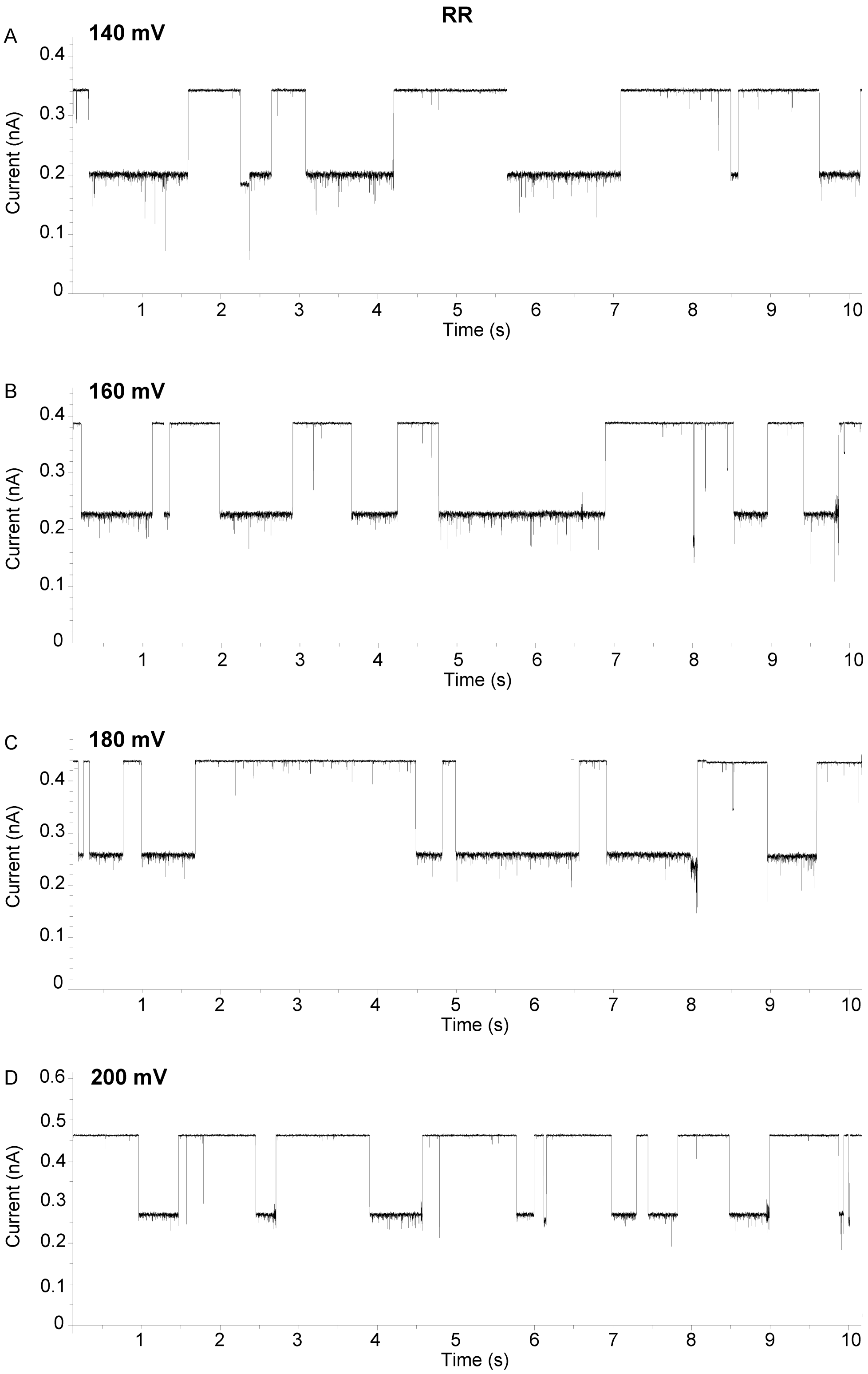


**Figure S2. Raw current traces of MC-RR.** Raw current traces at 140mV **(A)**, 160 mV **(B)**, 180 mV **(C)**, 200 mV **(D)** acquired in 4 M KCl, buffered with 10 mM Tris, 1.0 mM EDTA at pH 7.5; traces were filtered to 500 Hz. As the voltage increases, dwell times initially become longer (up to 160 mV), but decrease from 180 mV. Data extraction was performed as described in the Methods section.

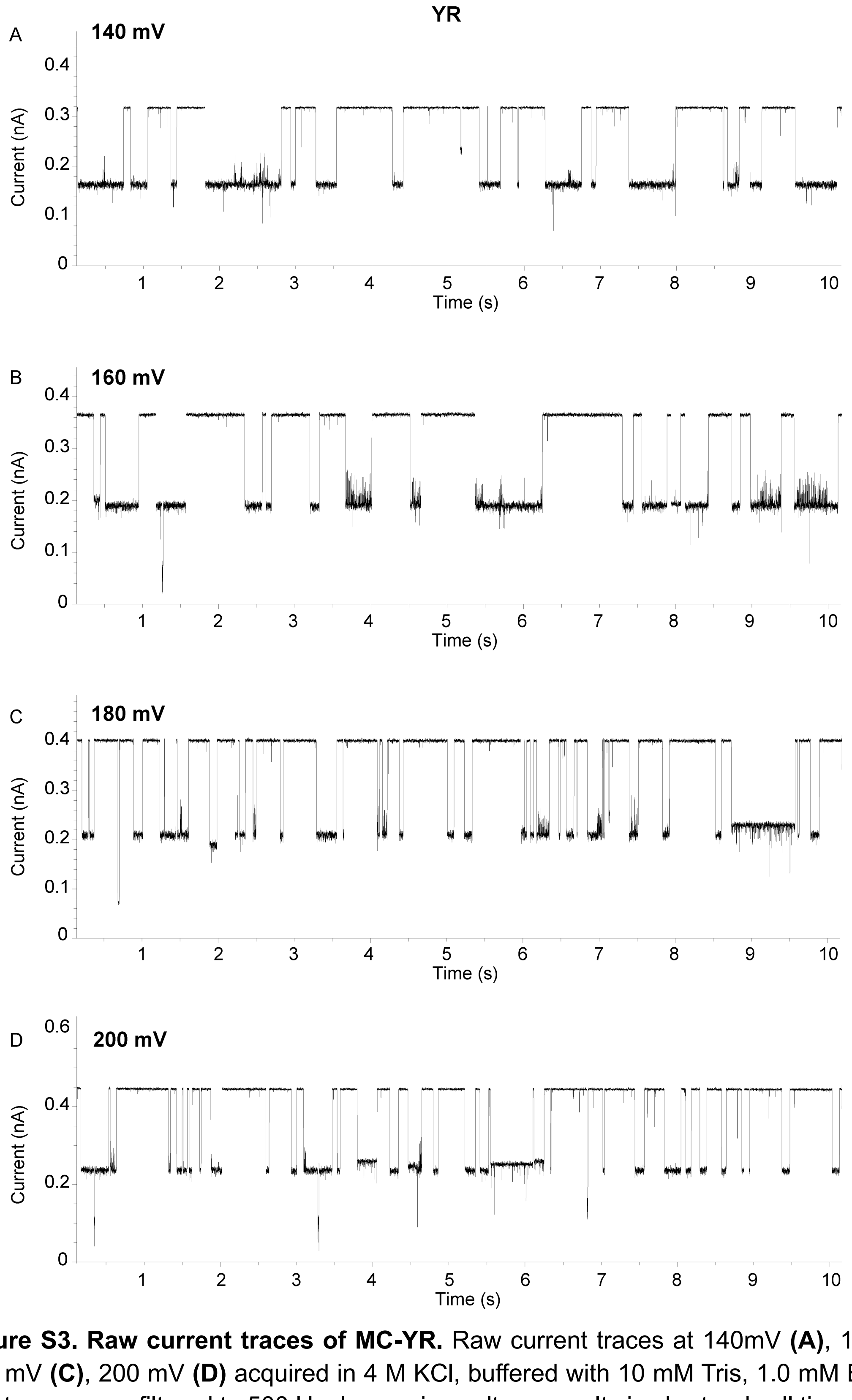


**Figure S3. Raw current traces of MC-YR.** Raw current traces at 140mV **(A)**, 160 mV **(B)**, 180 mV **(C)**, 200 mV **(D)** acquired in 4 M KCl, buffered with 10 mM Tris, 1.0 mM EDTA at pH 7.5; traces were filtered to 500 Hz. Increasing voltage results in shorter dwell times, as shown in the traces. Data extraction was performed as described in the Methods section.

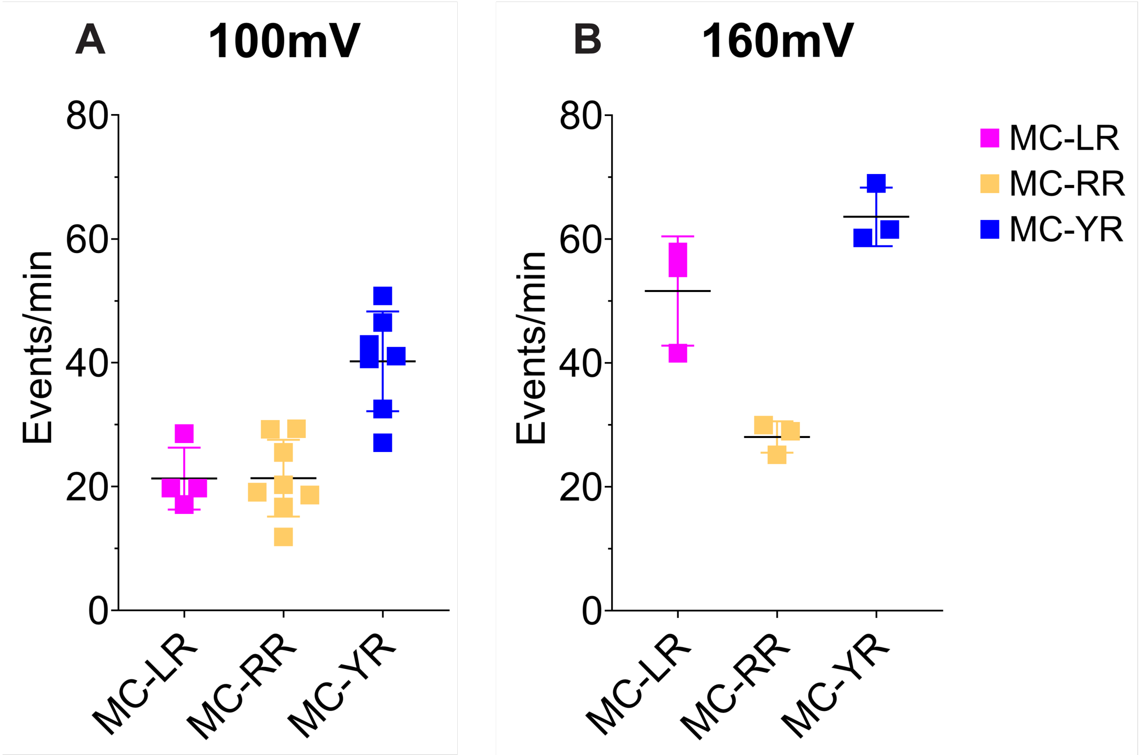


| Events per minute | MC-LR | MC-RR | MC-YR |
|---|---|---|---|
| **100 mV** | 21.3 ± 5.0 | 21.34 ± 6.19 | 40.22 ± 8.07 |
| **160 mV** | 51.62 ± 8.81 | 28.03 ± 2.53 | 63.58 ± 4.74 |

**Figure S4. Event rates analysis of MC-LR, -RR, -YR congeners in individual experiments at 100 mV and 160 mV.** Event rates (events/min) from single-pore recordings of each congener at 100 mV **(A)** and 160 mV **(B)**, where each square represents an individual pore. Mean values are shown below; 160 mV data were used to determine concentration ratios for uniform signal distribution in mixture experiments.

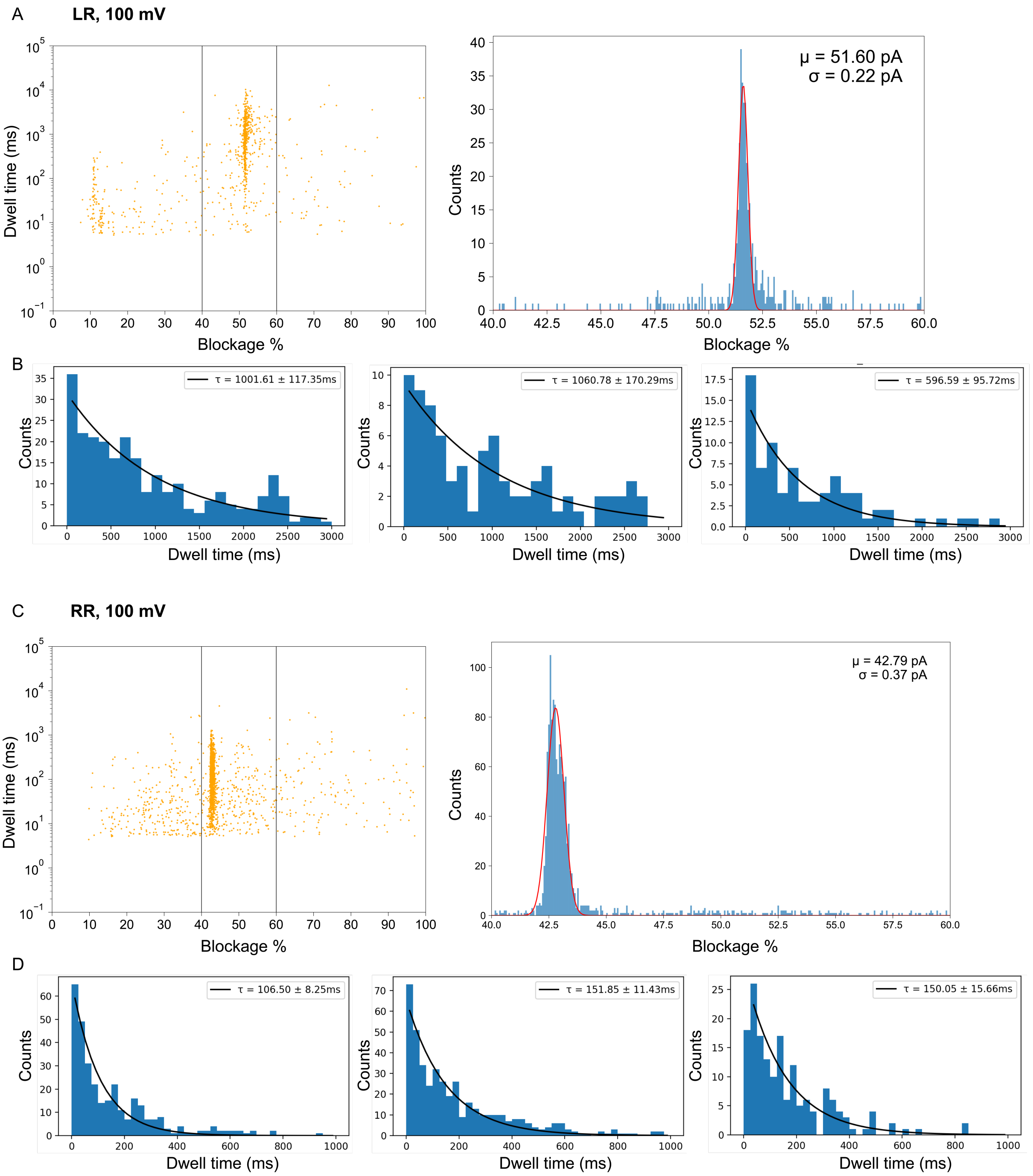

A
LR, 100 mV
Dwell time (ms)
Blockage %
μ = 51.60 pA
σ = 0.22 pA
Counts
B
τ = 1001.61 ± 117.35ms
τ = 1060.78 ± 170.29ms
τ = 596.59 ± 95.72ms
Dwell time (ms)
C
RR, 100 mV
μ = 42.79 pA
σ = 0.37 pA
D
τ = 106.50 ± 8.25ms
τ = 151.85 ± 11.43ms
τ = 150.05 ± 15.66ms

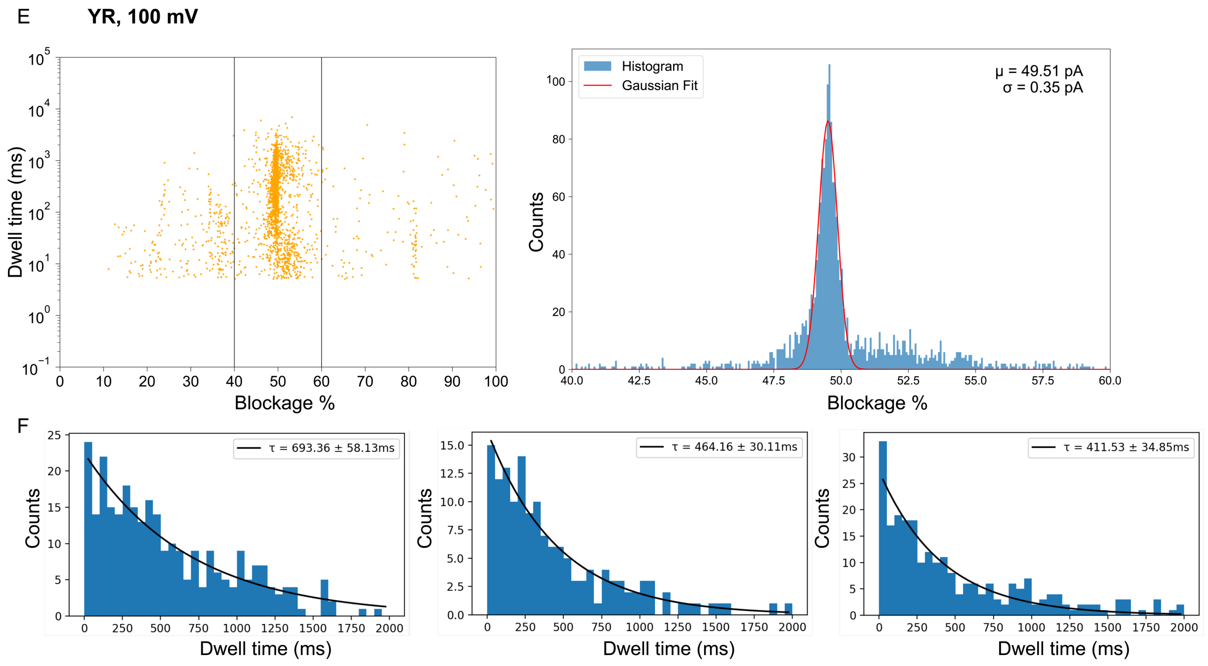


**Figure S5. Event selection of MC-LR, -RR, -YR at 100 mV. (A, C, E)** Scatterplots across at least 3 pore replicas produced by MC-LR, -RR, -YR interaction with WT AeL at 100 mV in individual experiments, respectively, event selection denoted by a box (left; 40 % - 60 %), and the current histograms produced by MC-LR, -RR, -YR at 100 mV (bin size = 300), respectively (right). **(B, D, F)** Exponential fitting (bin size = 20) of MC-LR, -RR, -YR dwell times, respectively, per single pore, with 3 representative individual pore replicas illustrated. Data extraction was performed as described in the Methods.

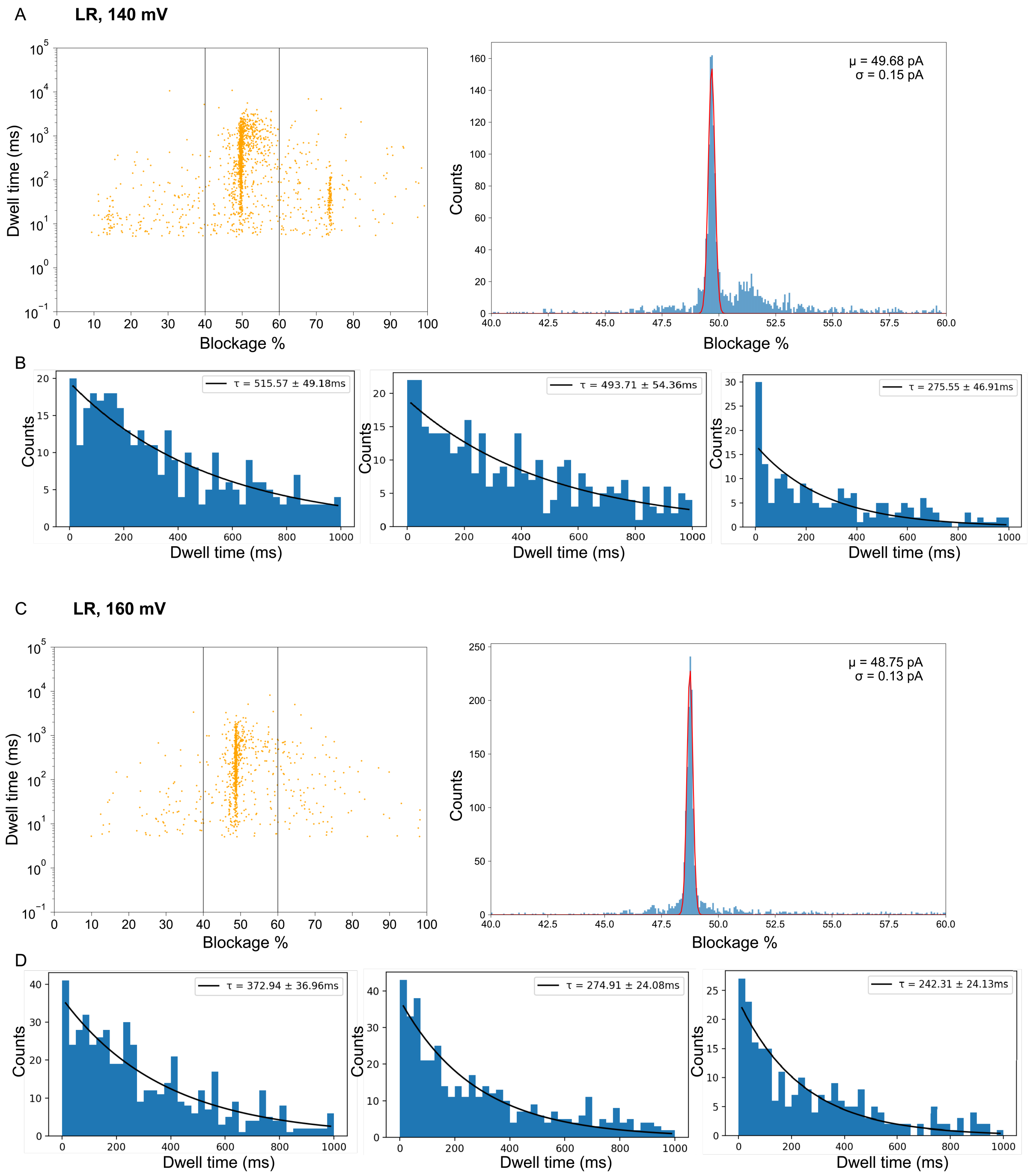
A
LR, 140 mV
Dwell time (ms)
Blockage %
Counts
μ = 49.68 pA
σ = 0.15 pA
B
τ = 515.57 ± 49.18ms
τ = 493.71 ± 54.36ms
τ = 275.55 ± 46.91ms
Dwell time (ms)
C
LR, 160 mV
μ = 48.75 pA
σ = 0.13 pA
D
τ = 372.94 ± 36.96ms
τ = 274.91 ± 24.08ms
τ = 242.31 ± 24.13ms

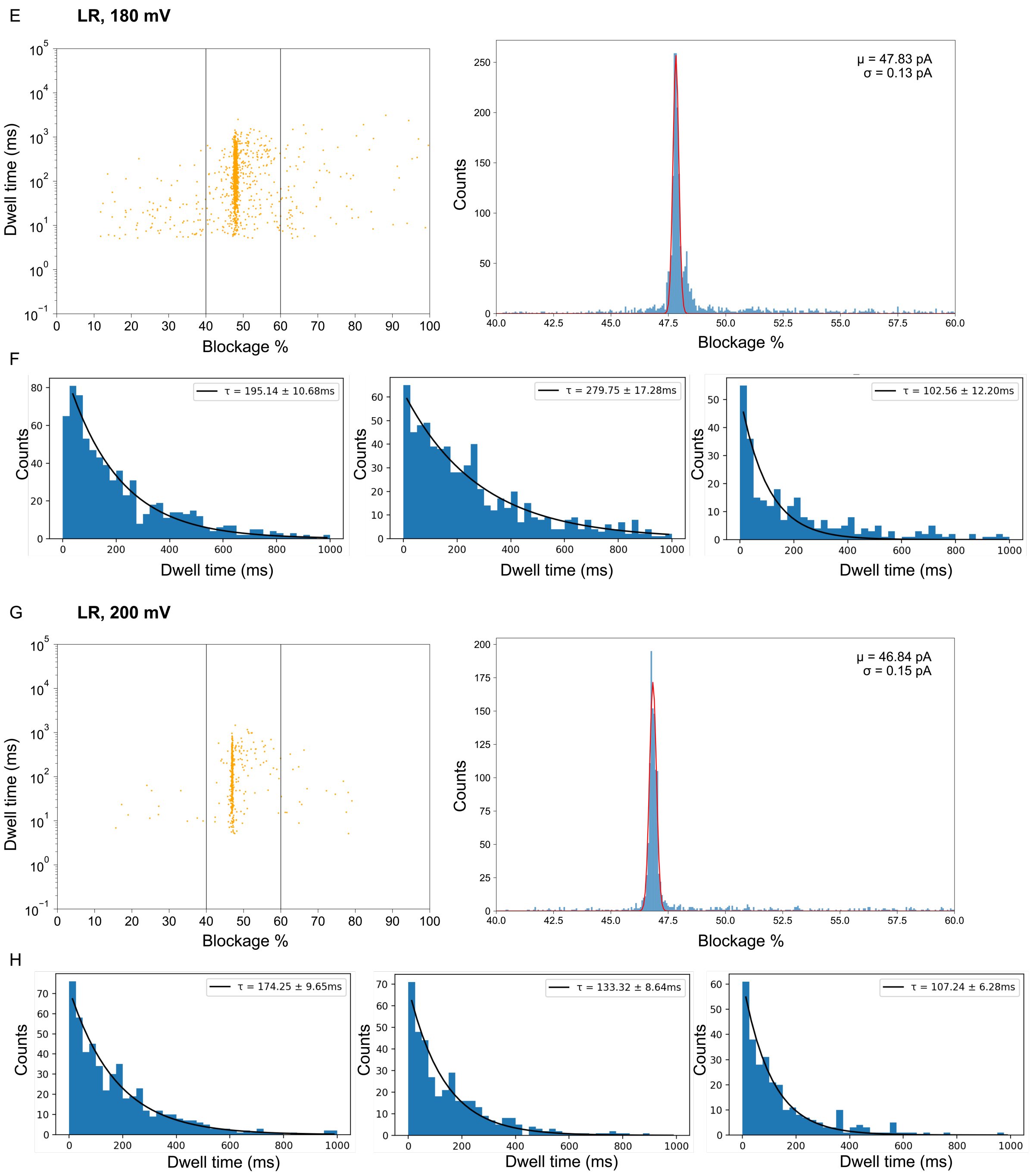


**Figure S6. Event selection of MC-LR at 140 mV, 160 mV, 180 mV, 200 mV. (A, C, E, G)** Scatterplots across at least 3 pore replicas, event selection denoted by a box (left; 40 % - 60 %), and the current histograms (bin size = 300) produced by MC-LR interaction with WT Ael at the specific voltage (right). **(B, D, F, H)** Exponential fitting (bin size = 20) of MC-LR dwell times per single pore, with 3 representative individual pore replicas illustrated. Data extraction was performed as described in the Methods.

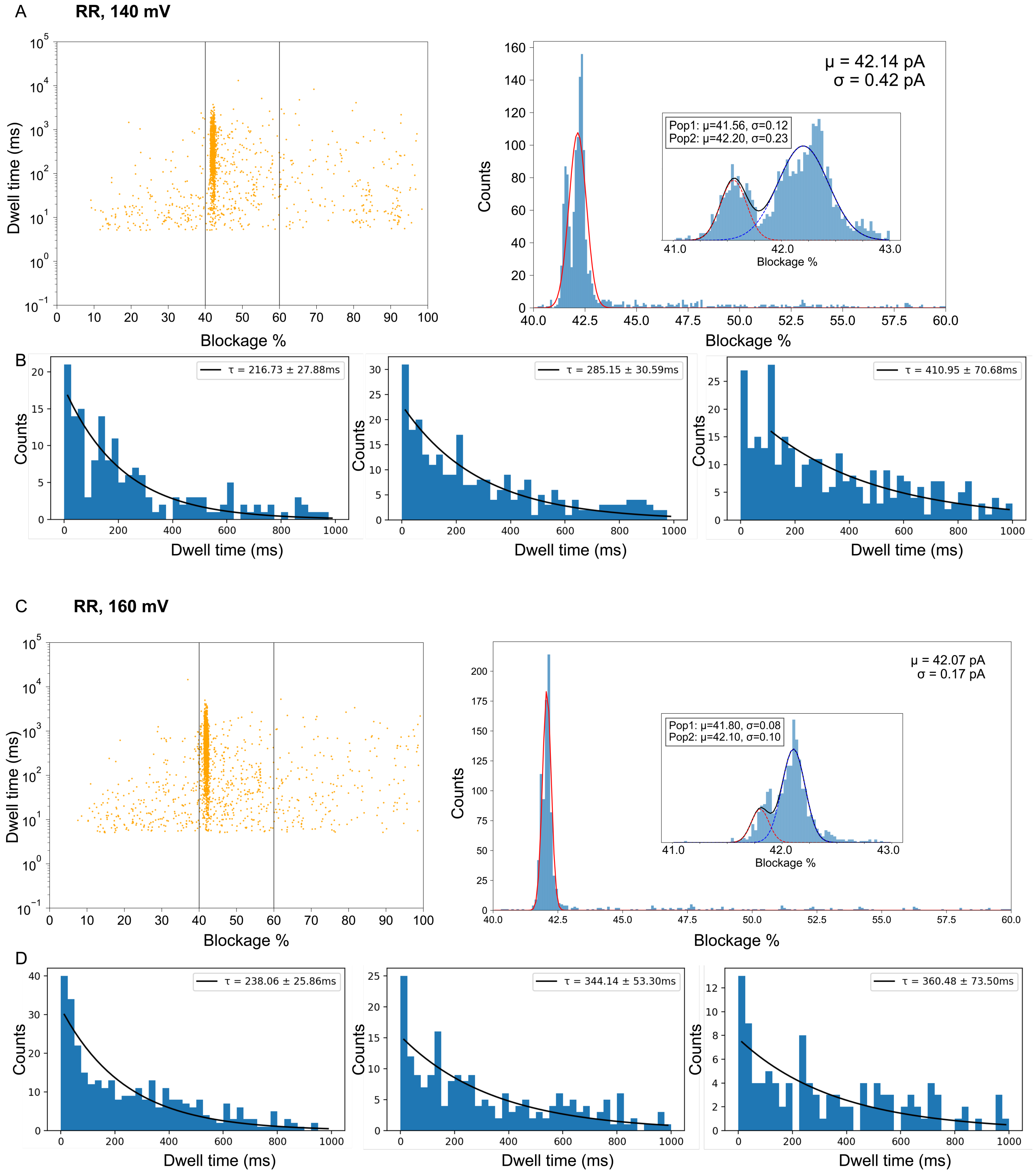
A
RR, 140 mV
Dwell time (ms)
Blockage %
Counts
μ = 42.14 pA
σ = 0.42 pA
Pop1: μ=41.56, σ=0.12
Pop2: μ=42.20, σ=0.23
Blockage %
B
τ = 216.73 ± 27.88ms
τ = 285.15 ± 30.59ms
τ = 410.95 ± 70.68ms
Counts
Dwell time (ms)
C
RR, 160 mV
Dwell time (ms)
Blockage %
Counts
μ = 42.07 pA
σ = 0.17 pA
Pop1: μ=41.80, σ=0.08
Pop2: μ=42.10, σ=0.10
Blockage %
D
τ = 238.06 ± 25.86ms
τ = 344.14 ± 53.30ms
τ = 360.48 ± 73.50ms
Counts
Dwell time (ms)

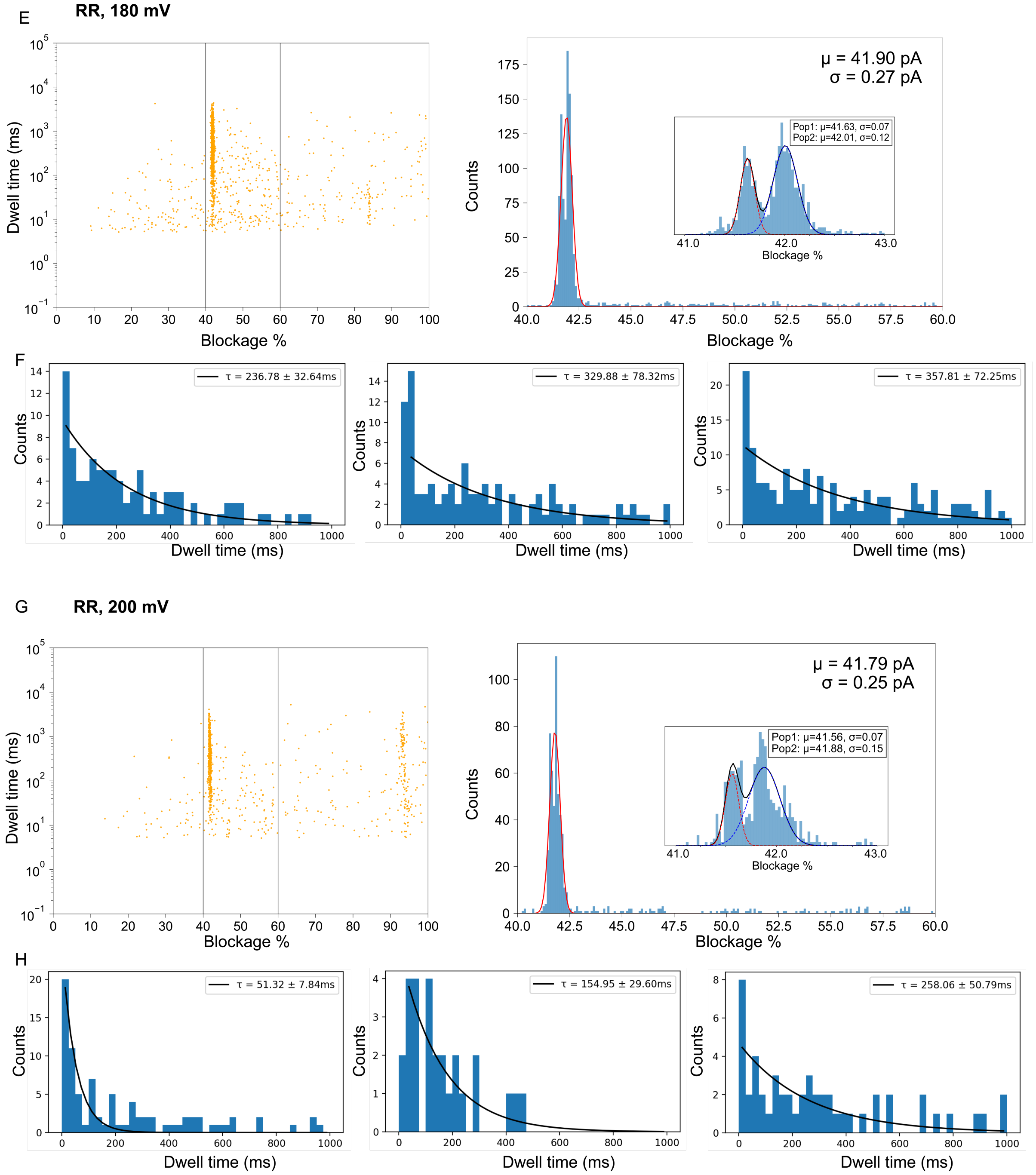


**Figure S7. Event selection of MC-RR at 140 mV, 160 mV, 180 mV, 200 mV. (A, C, E, G)** Scatterplots across at least 3 pore replicas, event selection denoted by a box (left; 40 % - 60 %), and the current histograms (bin size = 300 for single Gaussian fit; bin size = 75 for bimodal Gaussian fit) produced by MC-RR interaction with WT AeL at a specific voltage (right). **(B, D, F, H)** Exponential fitting (bin size = 20) of MC-RR dwell times per single pore, with 3 representative individual pore replicas illustrated. Data extraction was performed as described in the Methods.

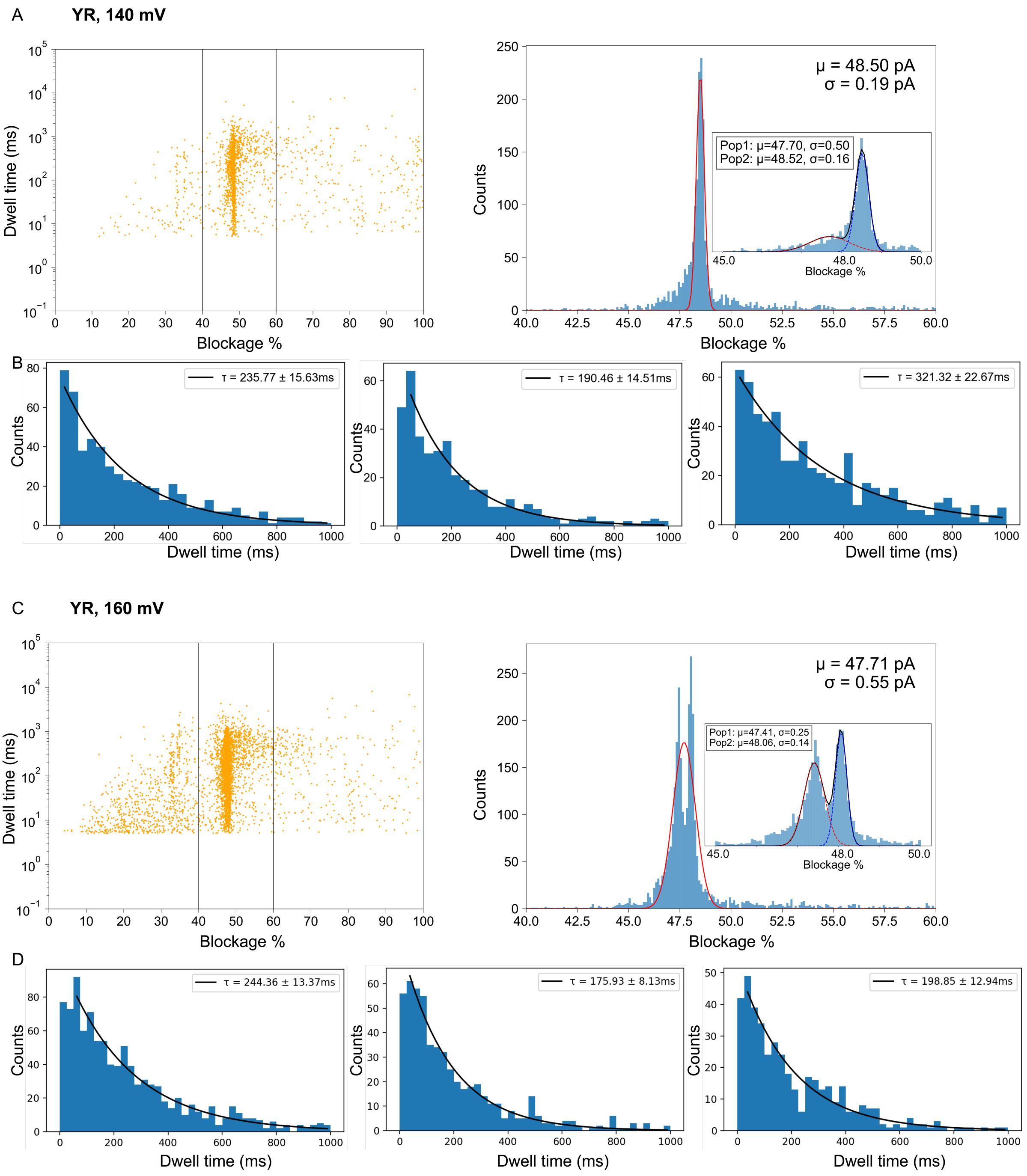
A
YR, 140 mV
Dwell time (ms)
Blockage %
Counts
μ = 48.50 pA
σ = 0.19 pA
Pop1: μ=47.70, σ=0.50
Pop2: μ=48.52, σ=0.16
B
τ = 235.77 ± 15.63ms
τ = 190.46 ± 14.51ms
τ = 321.32 ± 22.67ms
Dwell time (ms)
C
YR, 160 mV
μ = 47.71 pA
σ = 0.55 pA
Pop1: μ=47.41, σ=0.25
Pop2: μ=48.06, σ=0.14
D
τ = 244.36 ± 13.37ms
τ = 175.93 ± 8.13ms
τ = 198.85 ± 12.94ms

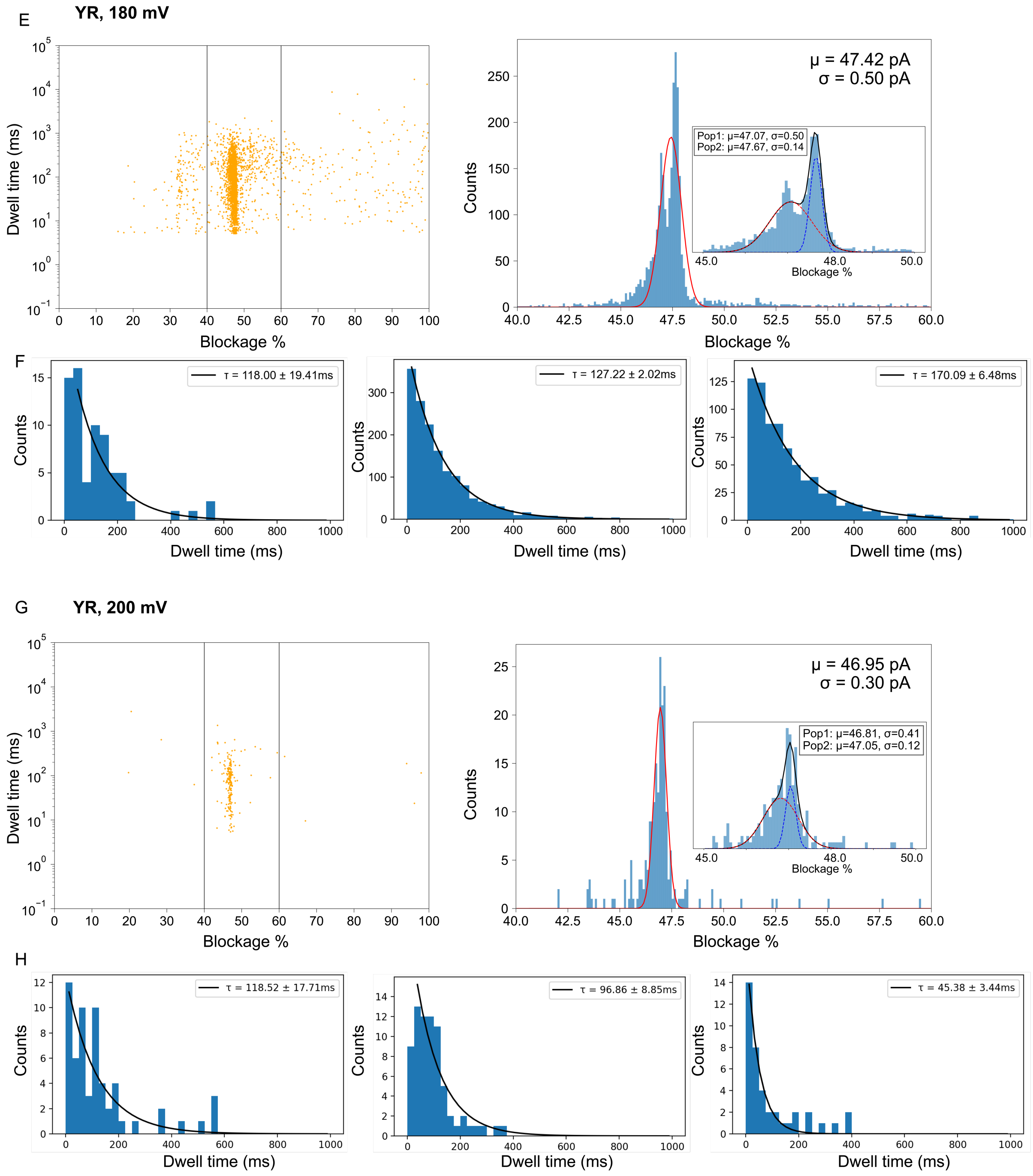


**Figure S8. Event selection of MC-YR at 140 mV, 160 mV, 180 mV, 200 mV. (A, C, E, G)** Scatterplots across at least 3 pore replicas, event selection denoted by a box (left; 40 % - 60 %), and the current histograms (bin size = 300 for single Gaussian fit; bin size = 75 for bimodal Gaussian fit) produced by MC-YR interaction with WT AeL at a specific voltage (right). **(B, D, F, H)** Exponential fitting (bin size = 20) of MC-YR dwell times per single pore, with 3 representative individual pore replicas illustrated. Data extraction was performed as in the Methods.

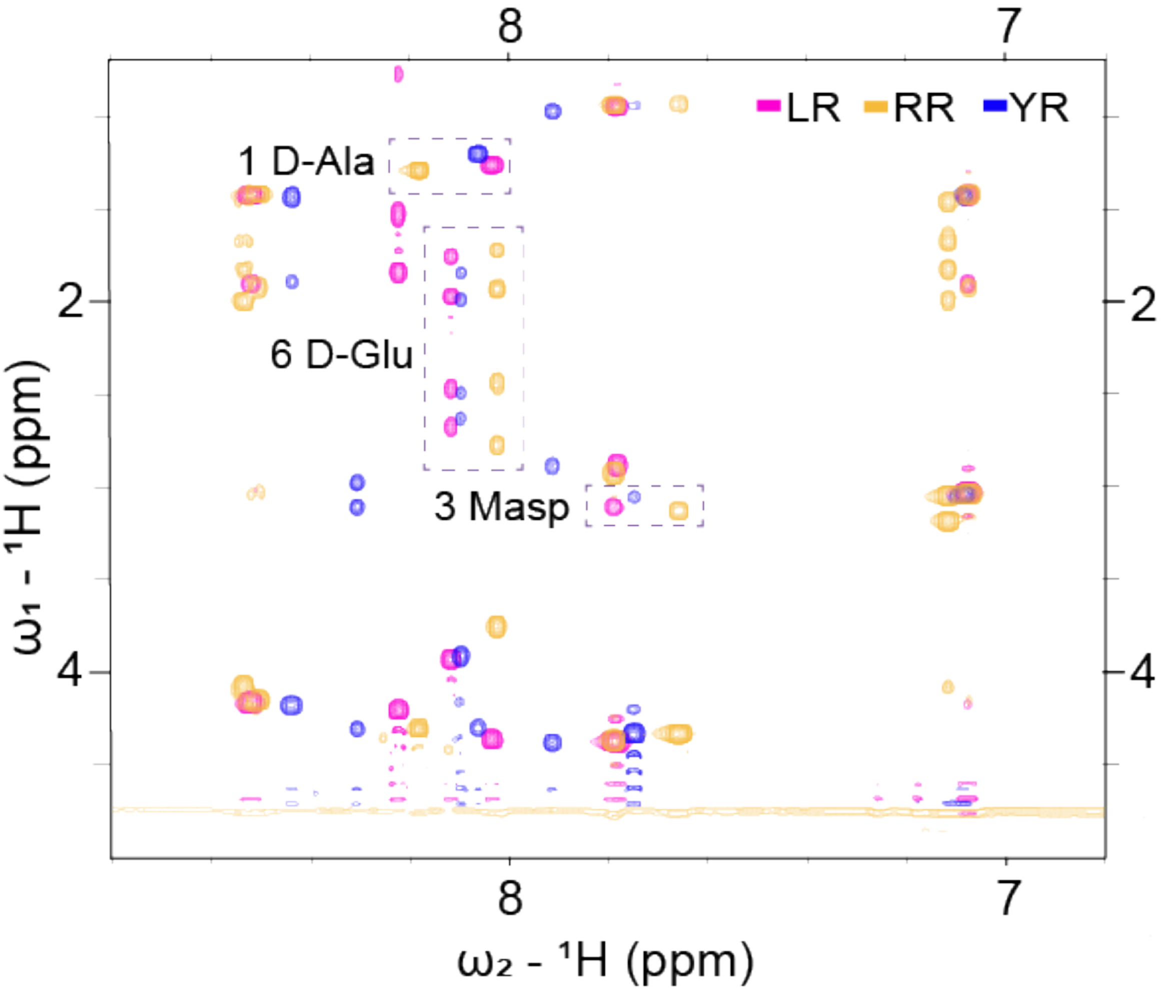


**Figure S9. TOCSY patterns emanating from HN groups in $^{1}$H,$^{1}$H TOCSY NMR spectra for MC-LR, -RR, -YR.** With boxes we highlight similarities and differences, labelled with assignments adapted to water conditions from published assignments[1–3] assisted with NOESY and $^{13}$C-HSQC spectra.

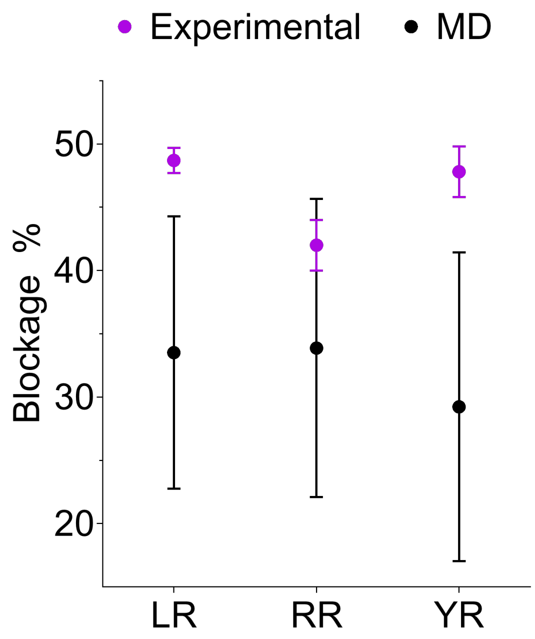


**Figure S10. Comparison of experimental and MD results yielding from MC-LR, -RR, -YR blockage of WT AeL at 160 mV.** MD simulations were performed as described in the Methods. Considering the significant standard deviation of the MD replicas, we conclude that MC-LR, -RR, and -YR behave the same within statistical significance, as determined by MD.

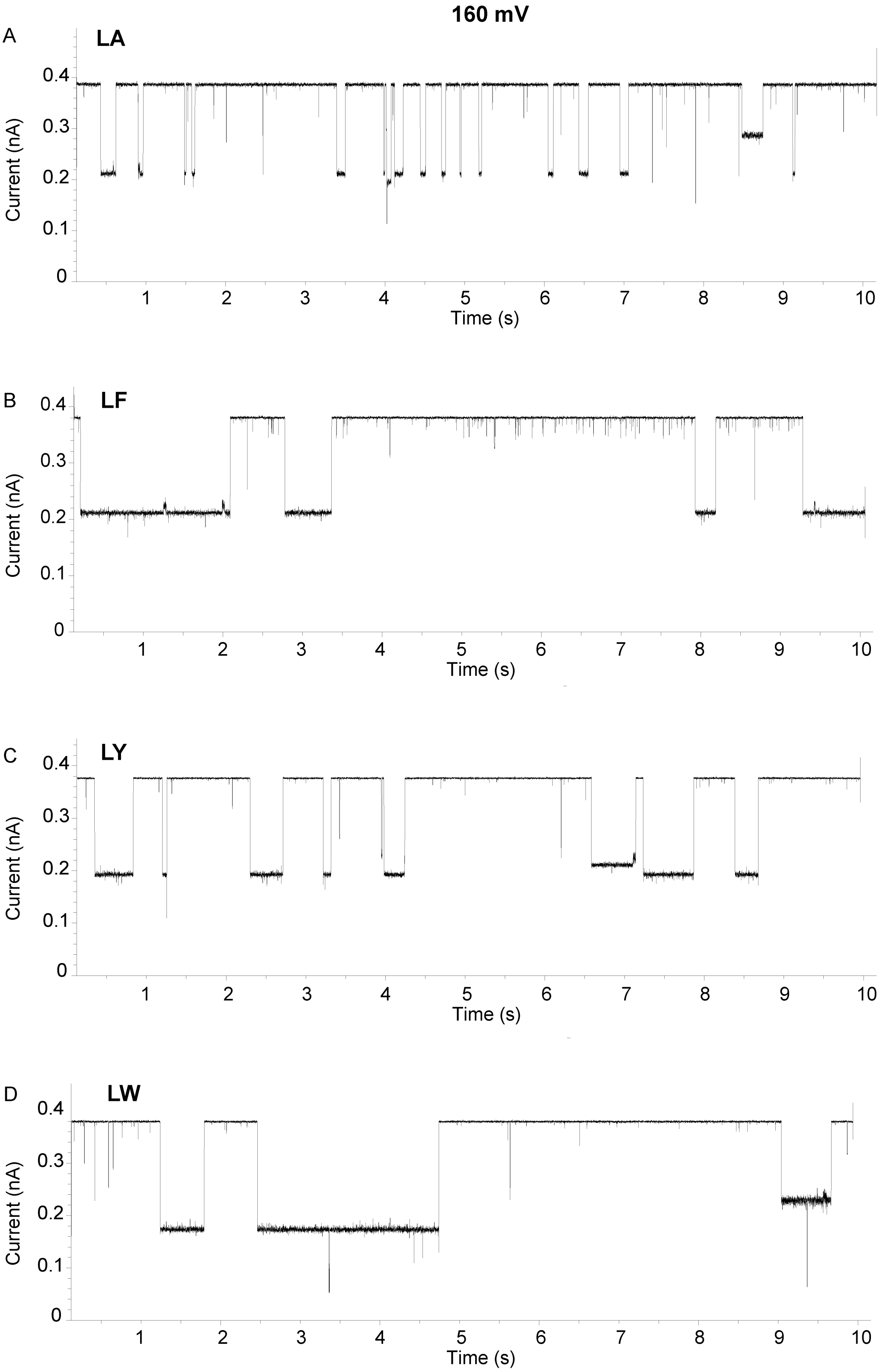


**Figure S11. Raw current traces of additional MCs at 160 mV.** Raw current traces of MC-LA **(A)**, -LF **(B)**, -LY **(C)**, -LW **(D)** acquired at 160 mV in 4 M KCl, buffered with 10 mM Tris, 1.0 mM EDTA at pH 7.5; traces were filtered to 500 Hz. Data extraction was performed as described in the Methods.

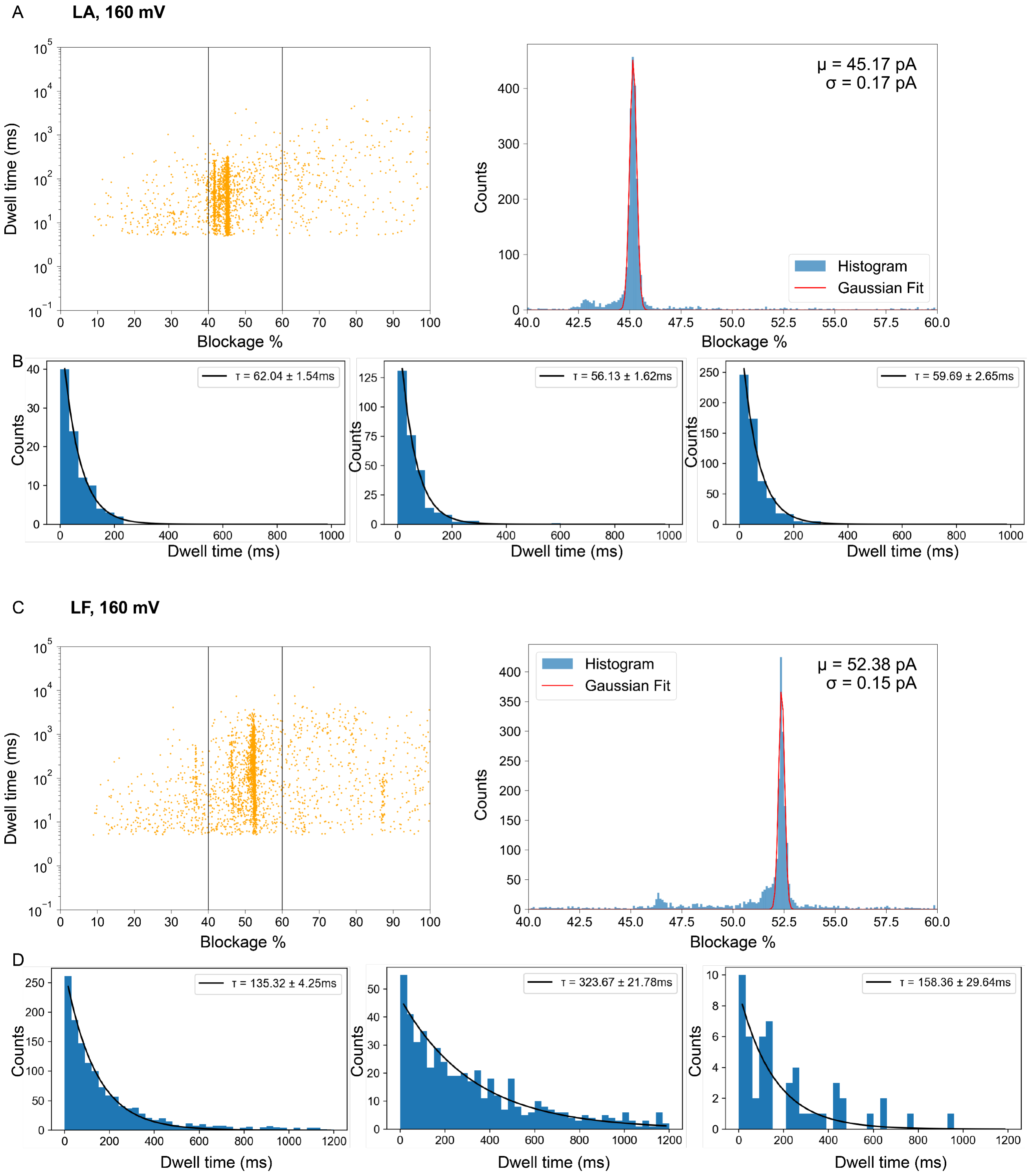
A
LA, 160 mV
Dwell time (ms)
Blockage %
μ = 45.17 pA
σ = 0.17 pA
Counts
Histogram
Gaussian Fit
B
τ = 62.04 ± 1.54ms
τ = 56.13 ± 1.62ms
τ = 59.69 ± 2.65ms
Counts
Dwell time (ms)
C
LF, 160 mV
Dwell time (ms)
Blockage %
μ = 52.38 pA
σ = 0.15 pA
Counts
Histogram
Gaussian Fit
D
τ = 135.32 ± 4.25ms
τ = 323.67 ± 21.78ms
τ = 158.36 ± 29.64ms
Counts
Dwell time (ms)

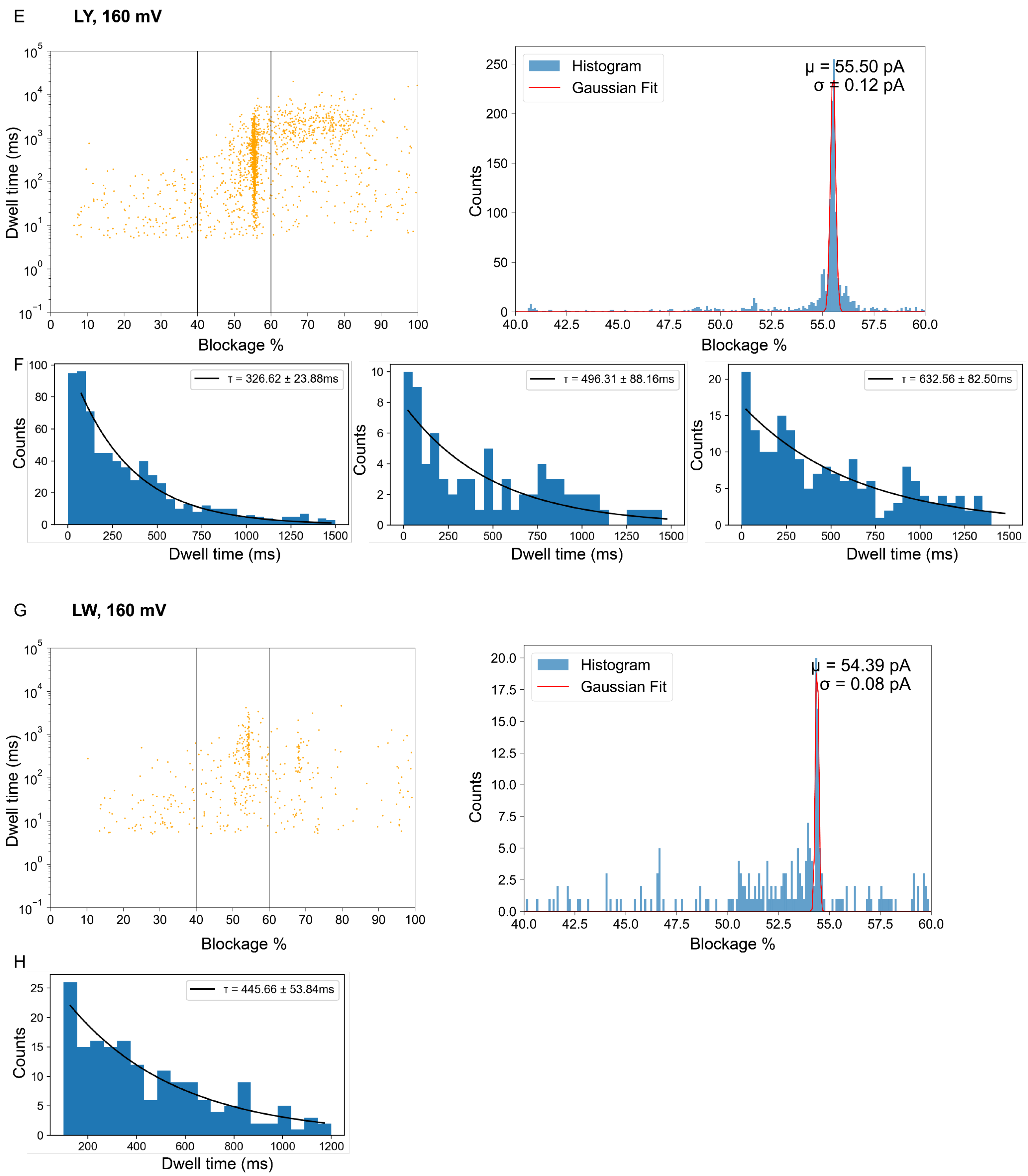


**Figure S12. Event selection of MC-LA, -LF, -LY, -LW at 160 mV. (A, C, E, G)** Scatterplots across at least 3 pore replicas, event selection denoted by a box (left; 40-60 %), and the histograms (bin size = 200) produced by the respective MC interaction with WT AeL (right). **(B, D, F)** Exponential fitting (bin size = 20) of the respective MC dwell times per single pore, with 3 representative individual pore replicas illustrated. **(H)** Exponential fitting of MC-LW dwell times was applied to pooled data from 3 pore replicas to compensate for low event counts per pore. Data extraction was performed as described in the Methods.

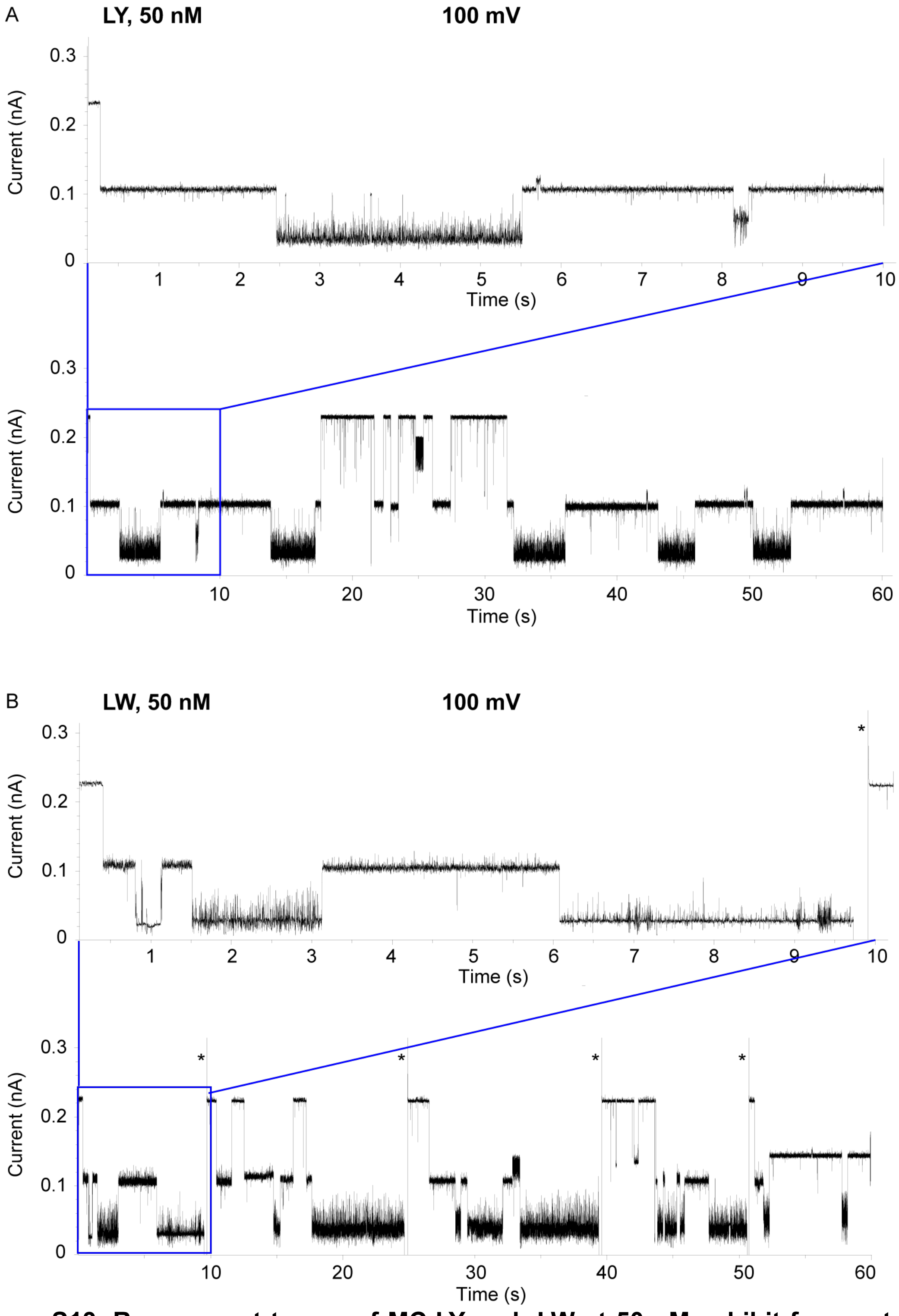


**Figure S13. Raw current traces of MC-LY and -LW at 50 nM exhibit frequent two-level blockade events.** Recordings were obtained at 100 mV in 4 M KCl, buffered with 10 mM Tris, 1.0 mM EDTA at pH 7.5; traces were filtered to 500 Hz. Panels **A** and **B** show traces for 50 nM MC-LY and 50 nM MC-LW, respectively, with a 10-second (top) and 60-second (bottom) time windows displayed in each panel. Stars (*) depict invert voltage to - 100 mV to unblock the pore, which frequently occurred with MC-LW under these conditions. The second (lower) level may indicate the presence of two MC molecules in the pore—one at the top and one at the bottom of the barrel. A transition from the second to the first level could reflect the upper MC displacing the lower one, initiating a repeating entry–exit cycle.

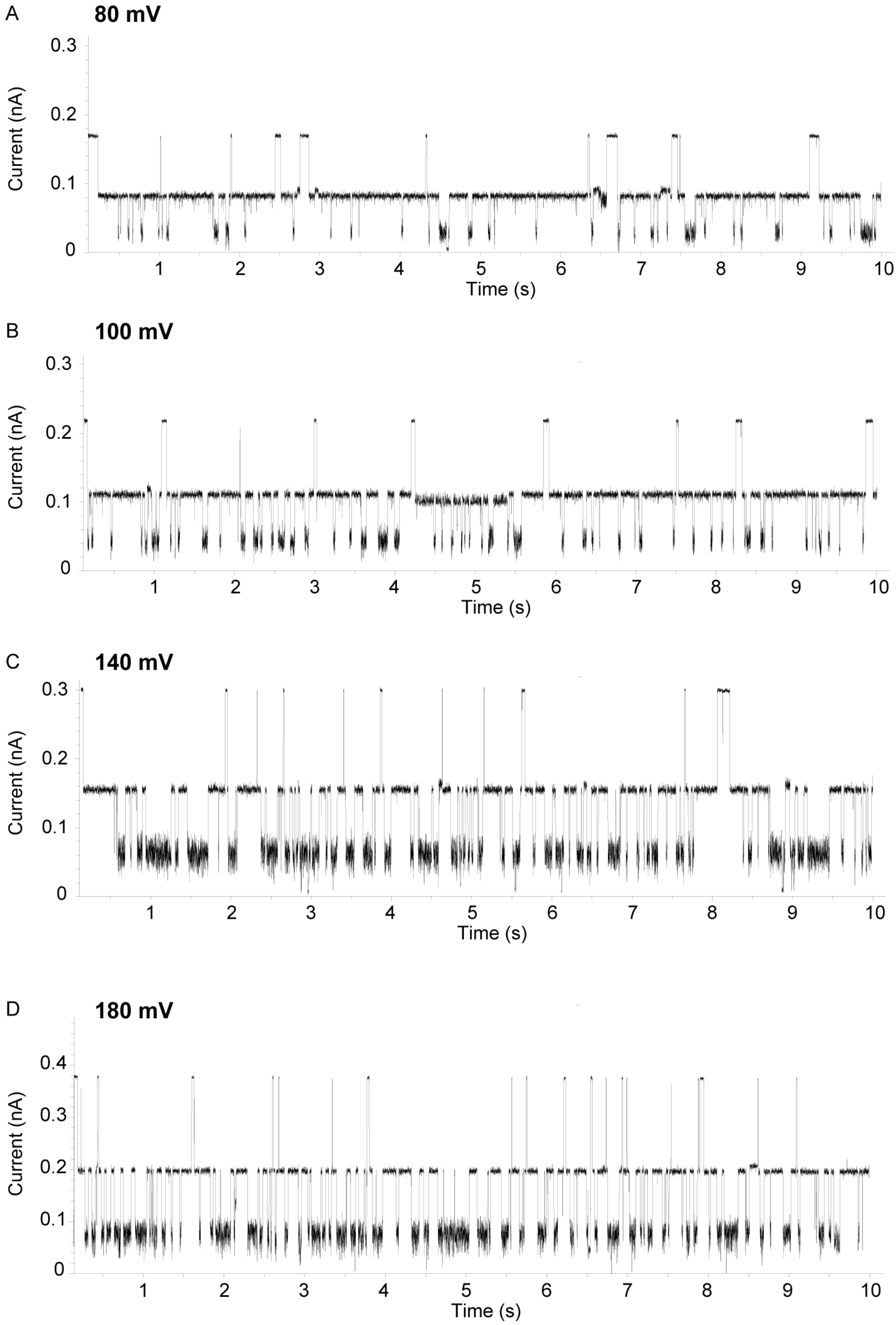


**Figure S14. Raw current traces of 2 μM MC-LR exhibit frequent two-level blockade events.** Recordings were obtained at 80 mV (**A**), 100 mV (**B**), 140 mV (**C**), 180 mV (**D**) in 4 M KCl, buffered with 10 mM Tris, 1.0 mM EDTA at pH 7.5; traces were filtered to 500 Hz. Voltage increase results in more frequent two-level blockades.

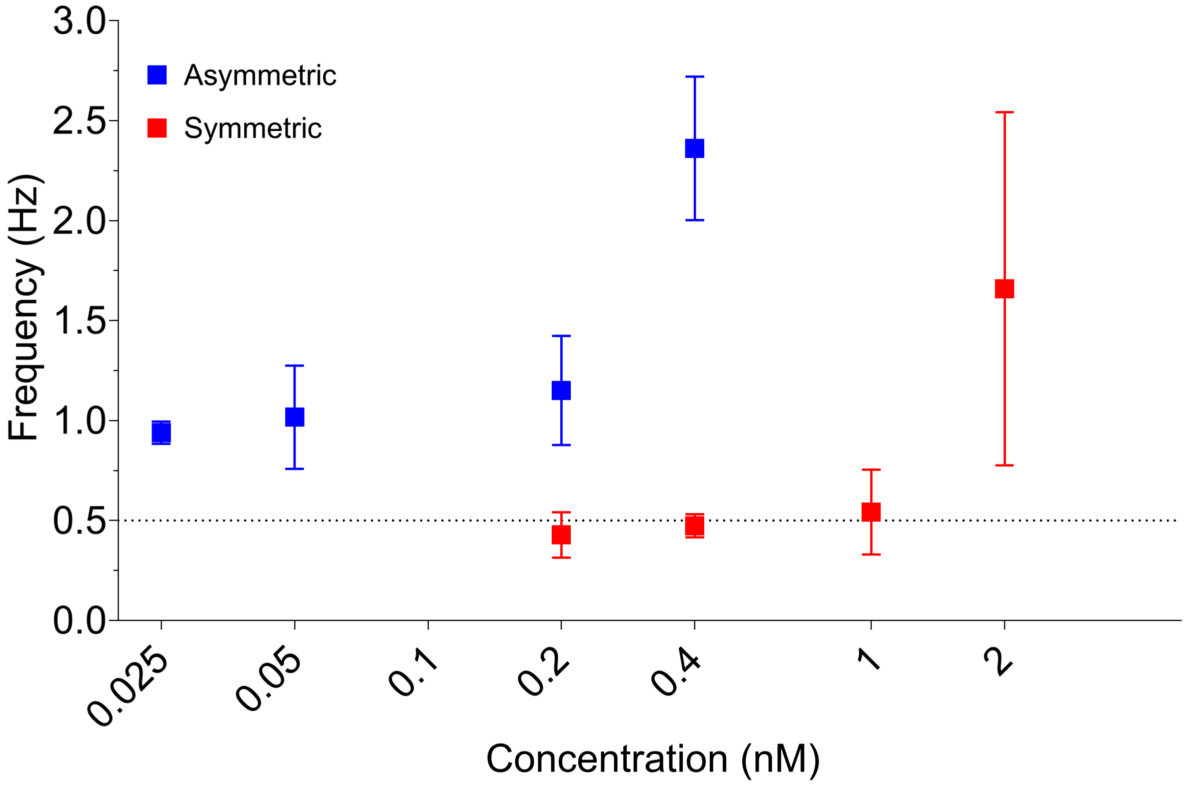


**Figure S15. Event frequency (Hz) of MC-LR in symmetric and asymmetric conditions.** Event frequency was calculated by an exponential fitting of inter-event time per pore (across 3 pore replicas), represented in Hz, across various concentrations in symmetric (red) and asymmetric (blue) conditions. Data collected at 100 mV under symmetric (4.0 M KCl) and asymmetric (0.15 M (*cis)* / 4.0 M (*trans)* KCl) conditions, both buffered with 10 mM Tris and 1.0 mM EDTA at pH 7.5.

### Supplementary Tables

**Table S1. Guideline values for MC-LR and exposure scenarios[4].**

| Toxin | Exposure | Value type (*µg/L*) |
|---|---|---|
| *Microcystin-LR* | Drinking-water, lifetime | 1 |
| *Microcystin-LR* | Drinking-water, short term | 12 |
| *Microcystin-LR* | Recreational | 24 |

**Table S2. Summary of physical and chemical properties of the tested MC congeners[5].** Weight and volume of the variable amino acid, weight and net charge of the respective microcystin congener.

| Amino acid X (position 2) | Weight (Da) | Volume (Å$^3$) | MCXR (Da) | MCXR Net charge |
|---|---|---|---|---|
| *L* | 131 | 166.7 | 995.2 | -1 |
| *R* | 174 | 173.4 | 1038.2 | 0 |
| *Y* | 181 | 193.6 | 1045.2 | -1 |

| Amino acid Z (position 4) | Weight (Da) | Volume (Å$^3$) | MCLZ (Da) | MCLZ Net charge |
|---|---|---|---|---|
| *A* | 89 | 88.6 | 910 | -2 |
| *F* | 165 | 189.9 | 986.2 | -2 |
| *Y* | 181 | 193.6 | 1002.2 | -2 |
| *W* | 204 | 227.8 | 1025.2 | -2 |

**Table S3. Analysis of $^3J_{HN\text{-}HA}$ coupling constants and $^{13}C\alpha$ and $^{13}C\beta$ chemical shifts for relevant MCLR and MCRR.**
Nres indicates residue number, with the canonical residue names shown on the first column. Cα and Cβ refer to the chemical shifts of $^{13}C$ nuclei in the MC backbones, akin to a small peptide (Cα on the cycle, Cβ first C atom in the sidechain). These chemical shifts reflect local conformational dynamics, interpreted as for peptide/protein chemical shifts by calculating $\Delta\delta_{C\alpha} - \Delta\delta_{C\beta}$, a calibration-insensitive probe that deviates from zero for structured regions, with higher values indicating stronger structuring. Similarly, $^3J_{HN\text{-}HA}$ reflects conformational averaging of the H-N-CA-HA dihedral, although here their differences are not too informative. In bold we highlight the key values discussed.

| | MC-LR | | | | | | | MC-RR | | | | | | | |
|---|---|---|---|---|---|---|---|---|---|---|---|---|---|---|---|
| **Residue** | **Nres** | $\delta_{C\alpha}$ | $\delta_{C\beta}$ | $\delta_{C\alpha}$-$\delta_{C\alpha rc}$* | $\delta_{C\beta}$-$\delta_{C\beta rc}$* | $\Delta\delta_{C\alpha}$- $\Delta\delta_{C\beta}$ | $^3J_{HN\text{-}HA}$ (Hz) | **Nres** | $\delta_{C\alpha}$ | $\delta_{C\beta}$ | $\delta_{C\alpha}$-$\delta_{C\alpha rc}$* | $\delta_{C\beta}$-$\delta_{C\beta rc}$* | $\Delta\delta_{C\alpha}$- $\Delta\delta_{C\beta}$ | $^3J_{HN\text{-}HA}$ (Hz) | **ΔJ (Hz)** |
| *Ala* | 1 | 49.54 | 15.92 | -2.961 | -3.08 | 0.121 | 6.92 | 1 | 50.01 | 15.94 | -2.491 | -3.06 | 0.57 | 6.44 | 0.48 |
| ***Leu/Arg*** | 2 | 53.59 | 39.22 | -2.114 | -2.68 | **0.569** | 7.27 | 2 | 55.56 | 28.2 | -0.743 | -2.1 | **1.361** | 6.62 | 0.65 |
| *Masp* | 3 | 55.99 | 40.87 | | | | 9.8 | 3 | 56.33 | 41.15 | | | | 9.98 | -0.2 |
| *Arg* | 4 | 51.68 | 27.22 | -4.624 | -3.08 | -1.54 | 9.8 | 4 | 51.59 | 27.33 | -4.708 | -2.97 | -1.74 | 9.92 | -0.1 |
| *ADDA* | 5 | 55.26 | 43.93 | | | | 9.2 | 5 | 55.5 | 43.82 | | | | 9.6 | -0.4 |
| ***Glu*** | 6 | 54.76 | 26.32 | -1.936 | -3.38 | **1.444** | 6.95 | 6 | 55.93 | 26.23 | -0.768 | -3.47 | **2.701** | 6.1 | 0.85 |
| *MDHA* | 7 | | 115.9 | | | | | 7 | | 115.1 | | | | | |

*$\delta_{C\alpha rc}$ and $\delta_{C\beta rc}$ are the random coil chemical shifts tabulated for each amino acid in [6].

**Table S4. Concentration-dependent ratio of MC-LR events.**
Each datapoint represents a single pore recording, where the ratio of MC-LR events is compared to the total events, plotted in **Fig. 5C**.

| | | | | | | | | | | | |
|---|---|---|---|---|---|---|---|---|---|---|---|
| **Asymmetric** | | | | | | | | | | | |
| Concentration (nM) | | | | | | | | | | | |
| 0.025 | 0.2988 | 0.1849 | 0.3333 | 0.2205 | 0.3333 | 0.1414 | 0.1 | 0.094 | 0.1011 | 0.1546 | 0.086 |
| 0.05 | 0.1573 | 0.1898 | 0.2857 | 0.1475 | 0.2061 | 0.1206 | 0.2098 | | | | |
| 0.1 | 0.4586 | 0.6316 | 0.644 | 0.5034 | 0.5921 | 0.4598 | | | | | |
| 0.2 | 0.7043 | 0.6087 | 0.5408 | 0.5088 | | | | | | | |
| 0.4 | 0.6018 | 0.584 | 0.5449 | 0.7021 | 0.6346 | | | | | | |
| **Symmetric** | | | | | | | | | | | |
| Concentration (nM) | | | | | | | | | | | |
| 0.2 | 0.0398 | 0.0235 | 0.0233 | 0.0606 | 0.0533 | 0.0824 | 0.0889 | | | | |
| 0.4 | 0.0414 | 0.0764 | 0.0874 | 0.0585 | 0.0294 | 0.0627 | 0.0906 | | | | |
| 1 | 0.2763 | 0.129 | 0.1026 | | | | | | | | |
| 2 | 0.6623 | 0.48 | 0.6565 | 0.4759 | | | | | | | |
| 20 | 0.7817 | 0.6763 | 0.7197 | 0.8077 | 0.7061 | 0.7061 | 0.7634 | | | | |
| 50 | 0.8571 | 0.7439 | 0.753 | | | | | | | | |
